# AAV-Txnip prolongs cone survival and vision in mouse models of retinitis pigmentosa

**DOI:** 10.1101/2021.01.27.428411

**Authors:** Yunlu Xue, Sean K. Wang, Parimal Rana, Emma R. West, Christin M. Hong, Helian Feng, David M. Wu, Constance L. Cepko

## Abstract

Retinitis pigmentosa (RP) is an inherited retinal disease, affecting >20 million people worldwide. Loss of daylight vision typically occurs due to the dysfunction/loss of cone photoreceptors, the cell type that initiates our color and high acuity vision. Currently, there is no effective treatment for RP, other than gene therapy for a limited number of specific disease genes. To develop a gene-agnostic therapy, we screened ≈20 genes for their ability to prolong cone photoreceptor survival *in vivo*. Here, we report an adeno-associated virus (AAV) vector expressing Txnip, which prolongs the survival of cone photoreceptors and improves visual acuity in RP mouse models. A Txnip allele, C247S, which blocks the association of Txnip with thioredoxin, provides an even greater benefit. Additionally, the rescue effect of Txnip depends on lactate dehydrogenase b (Ldhb), and correlates with the presence of healthier mitochondria, suggesting that Txnip saves RP cones by enhancing their lactate catabolism.

## Introduction

Retinitis pigmentosa (RP) is one of the most prevalent types of inherited retinal diseases, affecting approximately one in 3,000 people (Hartong et al., 2006). In RP, the rod photoreceptors, which initiate night vision, are primarily affected by the disease genes, and degenerate first. The degeneration of cones, the photoreceptors that initiate daylight, color and high acuity vision, then follows, which greatly impacts the quality of life. Currently, one therapy that holds great promise for RP is gene therapy using AAV (Maguire et al., 2019). This approach has proven successful for a small number of genes affecting a few disease families (Cehajic-Kapetanovic et al., 2020). However, due to the number and functional heterogeneity of RP disease genes (≈100 genes that primarily affect rods (https://sph.uth.edu/retnet/), gene therapy for each RP gene will be logistically and financially difficult. In addition, a considerable number of RP patients do not have an identified disease gene. A disease gene-agnostic treatment aimed at prolonging cone function/survival in the majority of RP patients could thus benefit many more patients. Given that the disease gene is typically not expressed in cones, and thus their death is due to non-autonomous mechanisms that may be in common across disease families, answers to the question of why cones die may provide an avenue to a widely applicable therapy for RP. To date, the suggested mechanisms of cone death include oxidative damage (Komeima et al., 2006; Wellard et al., 2005; Xiong et al., 2015), inflammation (Wang et al., 2020, 2019; Zhao et al., 2015), and a shortage of nutrients (Aït-Ali et al., 2015; Kanow et al., 2017; Punzo et al., 2012, 2009; Wang et al., 2016).

In 2009, we surveyed gene expression changes that occurred during retinal degeneration in four mouse models of RP (Punzo et al., 2009). Those data led us to suggest a model wherein cones starve and die due to a shortage of glucose, which is typically used for energy and anabolic needs in photoreceptors via glycolysis. Evidence of this “glucose shortage hypothesis” was subsequently provided by orthogonal approaches from other groups (Aït-Ali et al., 2015; Wang et al., 2016). These studies have inspired us to test ≈20 genes that might affect the uptake and/or utilization of glucose by cones *in vivo* in three mouse models of RP (Supplementary Table 1). Only one gene, *txnip*, had a beneficial effect, prolonging cone survival and visual acuity in these models. *Txnip* encodes an α-arrestin family member protein with multiple functions, including binding to thioredoxin (Junn et al., 2000; Nishiyama et al., 1999), facilitating removal of the glucose transporter 1 (Glut1), from the cell membrane (Wu et al., 2013), and promoting the use of non-glucose fuels (DeBalsi et al., 2014). Because α-arrestins are structurally distinct from the visual or β-arrestins such as Arr3, Txnip is unlikely to bind to opsins or to participate in phototransduction (Hwang et al., 2014; Kang et al., 2015; Puca and Brou, 2014). We tested a number of *txnip* alleles and found that one allele, C247S, which blocks the association of Txnip with thioredoxin (Patwari et al., 2009), provided the greatest benefit. Investigation of the mechanism of Txnip rescue revealed that it required lactate dehydrogenase b (Ldhb), which catalyzes the conversion of lactate to pyruvate. Imaging of metabolic reporters demonstrated an enhanced cytosolic ATP:ADP ratio when the retina was placed in lactate medium. Moreover, mitochondria appeared to be healthier as a result of Txnip addition, but this improvement was not sufficient for cone rescue.

The above observations led to a model wherein Txnip shifts cones from their normal reliance on glucose to enhanced utilization of lactate, as well as marked improvement in mitochondrial structure and function. Analysis of the rescue activity of several additional genes predicted to affect glycolysis, provided support for this model. Finally, as our goal is to rescue cones that suffer not only from metabolic challenges, but also from inflammation and oxidative damage, we tested Txnip in combination with anti-inflammatory and anti-oxidative damage genes, and found additive benefits for cones. These treatments may benefit cones not only in RP, but also in other ocular diseases where similar environmental stresses are present, such as in age related macular degeneration (AMD).

## Results

### Txnip prolongs RP cone survival and visual acuity

We delivered genes that might address a glucose shortage and/or mismanagement of metabolism in a potentially glucose-limited environment. To this end, twelve AAV vectors were constructed to test genes singly or in combination for an initial screen (Extended Data Fig. 1e). Subsequently, an additional set of AAV vectors were made based upon the initial screen results, as well as other rationales, to total 20 genes tested in all (Supplementary Table 1). Most of these vectors carried genes to augment the utilization of glucose, such as hexokinases (Hk1 and Hk2), phosphofructokinase (Pfkm) and pyruvate kinase (Pkm1 and Pkm2). Each AAV vector used a cone-specific promoter, which was previously found to be non-toxic at the doses used in this study (Xiong et al., 2019). An initial screen was carried out in *rd1* mice, which harbor a null allele in the rod-specific gene, *Pde6b*. This strain has a rapid loss of rods, followed by cone death. The vectors were subretinally injected into the eyes of neonatal *rd1* mice, in combination with a vector using the human red opsin (RedO) promoter to express a histone 2B-GFP fusion protein (AAV-RedO-H2BGFP). The H2BGFP provides a very bright cone-specific nuclear labelling, enabling automated quantification. As a control, eyes were injected with AAV-RedO-H2BGFP alone. *Rd1* cones begin to die at ≈postnatal day 20 (P20), when almost all rods have died (Extended Data Fig. 1a). The number of *rd1* cones was quantified by counting the H2BGFP+ cells using a custom-made MATLAB program (Fig. 1a and Extended Data Fig. 1c). Only cones within the central region of the retina were counted, since RP cones in the periphery die much later (Hartong et al., 2006; Punzo et al., 2009). Among the twelve tested vectors, and six of their combinations, we found that only Txnip led to an increase in P50 *rd1* cones. The effects were likely on cone survival, as it did not change the number of cones at P20 prior to their death (Fig. 1a,b, and Extended Data Fig. 1c,e). The level of Txnip rescue in P50 *rd1* cones was comparable to using AAV with a CMV promoter to express a transcription factor, Nrf2, that regulates anti-oxidation pathways and reduces inflammation as we found previously (Xiong et al., 2015) (Extended Data Fig. 1e). One combination led to a reduction in cone survival, that of Hk1 plus Pfkm (Extended Data Fig. 1e).

**Fig. 1:**
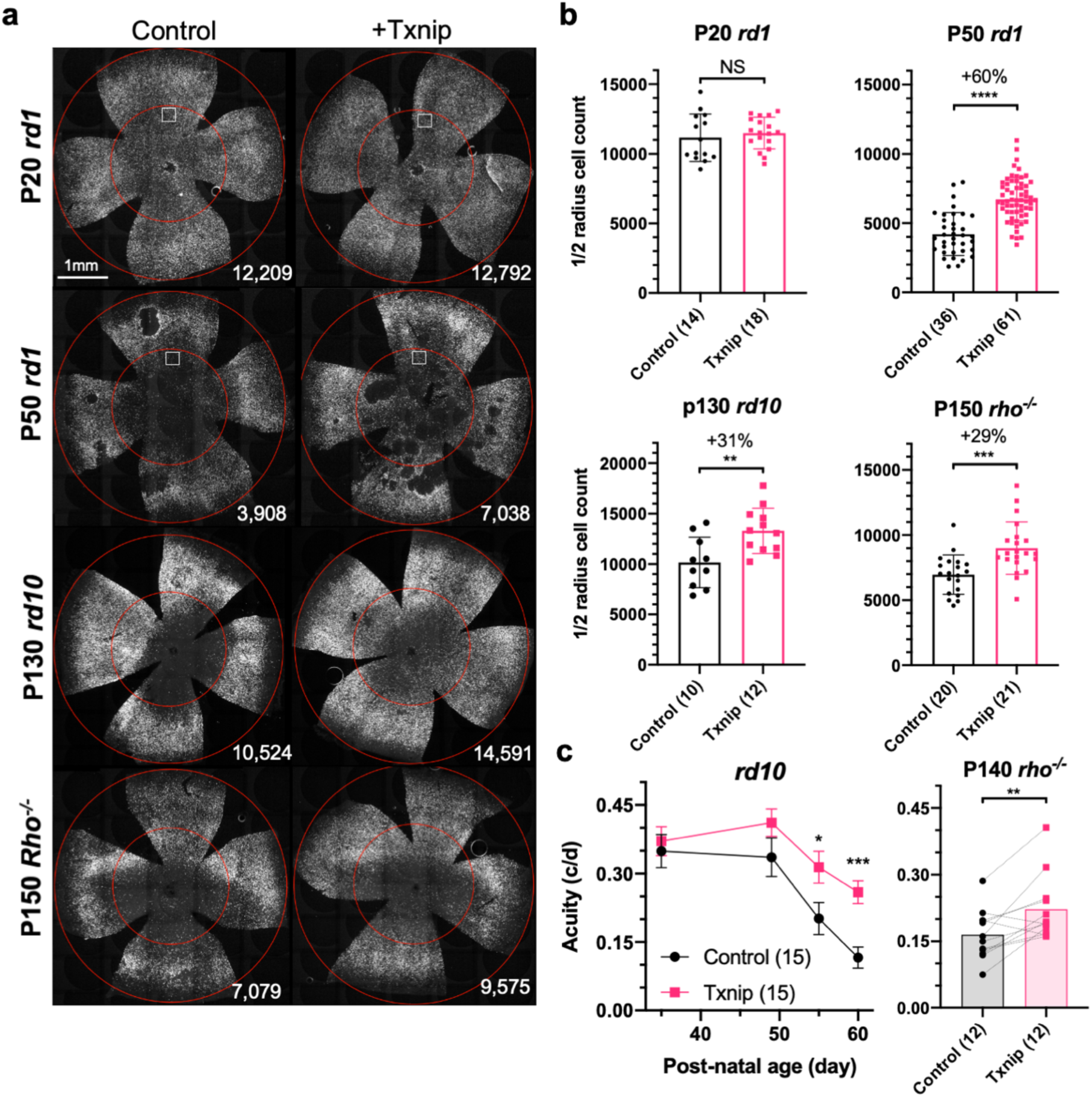
Txnip enhances cone survival and delays the deterioration of cone-mediated vision in RP mice. **a.** Representative images from P20 and P50 *rd1*, P130 *rd10* and P150 *rho*^-/-^ flat-mounted retinas, in which cones are labeled with H2BGFP, treated with Txnip or control (i.e. H2BGFP and vehicle only, same applies to all other figures). The outer circle was drawn to mark the full extent of the retina, and the inner circle was drawn by using half of the radius of the outer circle. The small box in the top four panels are zoomed in with pixels recognized as cones by a MATLAB automated-counting program (Extended Data Fig.1c). The number at lower right corner is the count of cells within the half radius of each image (same applies to all other figures). **b.** Quantification of H2BGFP-positive cones within the inner half of the retina at different groups. Error bar: standard deviation. The number in round brackets “()” indicates the sample size, i.e. the number of eyes/retinas within each group (same applies to all other figures). **c.** Visual acuity of *rd10* and P140 *rho*^-/-^ mice with Txnip or control treatment in each eye measured with optomotor assays. Error bar: SEM. NS: not significant, *p* > 0.05. * *p* < 0.05. ** *p* < 0.01. *** *p* < 0.001. **** *p* < or << 0.0001.

Our initial screen used the RedO promoter to drive Txnip expression. To evaluate a different cone-specific promoter, Txnip also was tested using a newly described cone-specific promoter, SynPVI (Jüttner et al., 2019). This promoter also led to prolonged cone survival (Extended Data Fig. 1e). To explore whether Txnip gene therapy is effective beyond *rd1*, it was tested in *rd10* mice, which carry a missense *Pde6b* mutation, and in *rho*^-/-^ mice, which carry a null allele in a rod-specific protein, rhodopsin. Prolonged cone survival was observed in both strains (Fig. 1a,b). To determine if Txnip-treated mice sustained greater visual acuity than control RP mice, an optomotor assay was used (Prusky et al., 2004). Under conditions that simulated daylight, Txnip treated eyes showed enhanced visual acuity compared to the control contralateral eyes in *rd10* and *rho*^-/-^ mice (Fig. 1c). Txnip also was evaluated for effects on cones in wildtype (wt) mice, using PNA staining, which stains the cone-specific extracellular matrix and reflects cone health. The approximate number and morphology of Txnip-treated cones appeared normal by this assay (Extended Data Fig. 1d).

### Evaluation of *txnip* alleles for cone survival

Previous studies of Txnip provided a number of alleles that could potentially lead to a more effective cone rescue by Txnip, and/or provide some insight into which of the Txnip functions are required for enhancing cone survival. A C247S mutation has been shown to block Txnip’s inhibitory interaction with thioredoxin (Patwari et al., 2009), which is an important component of a cell’s ability to fight oxidative damage via thiol groups (Junn et al., 2000; Nishinaka et al., 2001; Nishiyama et al., 1999). If cone rescue by Txnip required this function, the C247S allele should be less potent for cone rescue. Alternatively, if loss of thioredoxin binding freed Txnip for its other functions, and made more thioredoxin available for oxidative damage control, this allele might more effectively promote cone survival. The C247S clearly provided more robust cone rescue than wildtype (wt) Txnip in all three RP mouse strains (Fig. 2a,b and Extended Data Fig. 2a,b). These results indicate that the therapeutic effect of Txnip is not based on the inhibitory interaction with thioredoxins. This finding is in keeping with previous work which showed that anti-oxidation strategies promoted cone survival in RP mice (Komeima et al., 2006; Wu et al., n.d.; Xiong et al., 2015). An additional mutant, S308A, which loses an AMPK/Akt-phosphorylation site on Txnip (Waldhart et al., 2017; Wu et al., 2013), was tested in the context of wt Txnip and in the context of the C247S allele. The S308A allele did not benefit cone survival in either context (Fig. 2a,b). In addition, the S308A allele was assayed for negative effects on cones by an assessment of *rd1* cone number prior to P20, i.e. before the onset of cone death (Extended Data Fig. 2c). It did not reduce the cone number at this early timepoint, indicating that Txnip.S308A was not toxic to cones. This finding suggests that the S308 residue is critical for the therapeutic function of Txnip, through an unclear mechanism. One additional allele, LL351&352AA, was tested in the context of C247S. This allele eliminates a clathrin-binding site, and thus hampers Txnip’s ability to remove Glut1 from cell surface through clathrin-coated pits (Wu et al., 2013). Txnip.C247S.LL351&352AA could still delay RP cone death compared to the control (Fig. 2b), suggesting that the therapeutic effect of Txnip was unlikely to be only through the removal of Glut1 from the cell surface. To further explore the role of Glut1, an shRNA to *slc2a1*, which encodes Glut1, was tested. It did not prolong RP cone survival (Extended Data Fig. 2d). The slight decrease of Txnip.C247S.LL351&352AA in cone rescue compared to Txnip.C247S might be due to other, currently unknown effects of LL351&352, or a less specific effect, e.g. a protein conformational change.

**Fig. 2:**
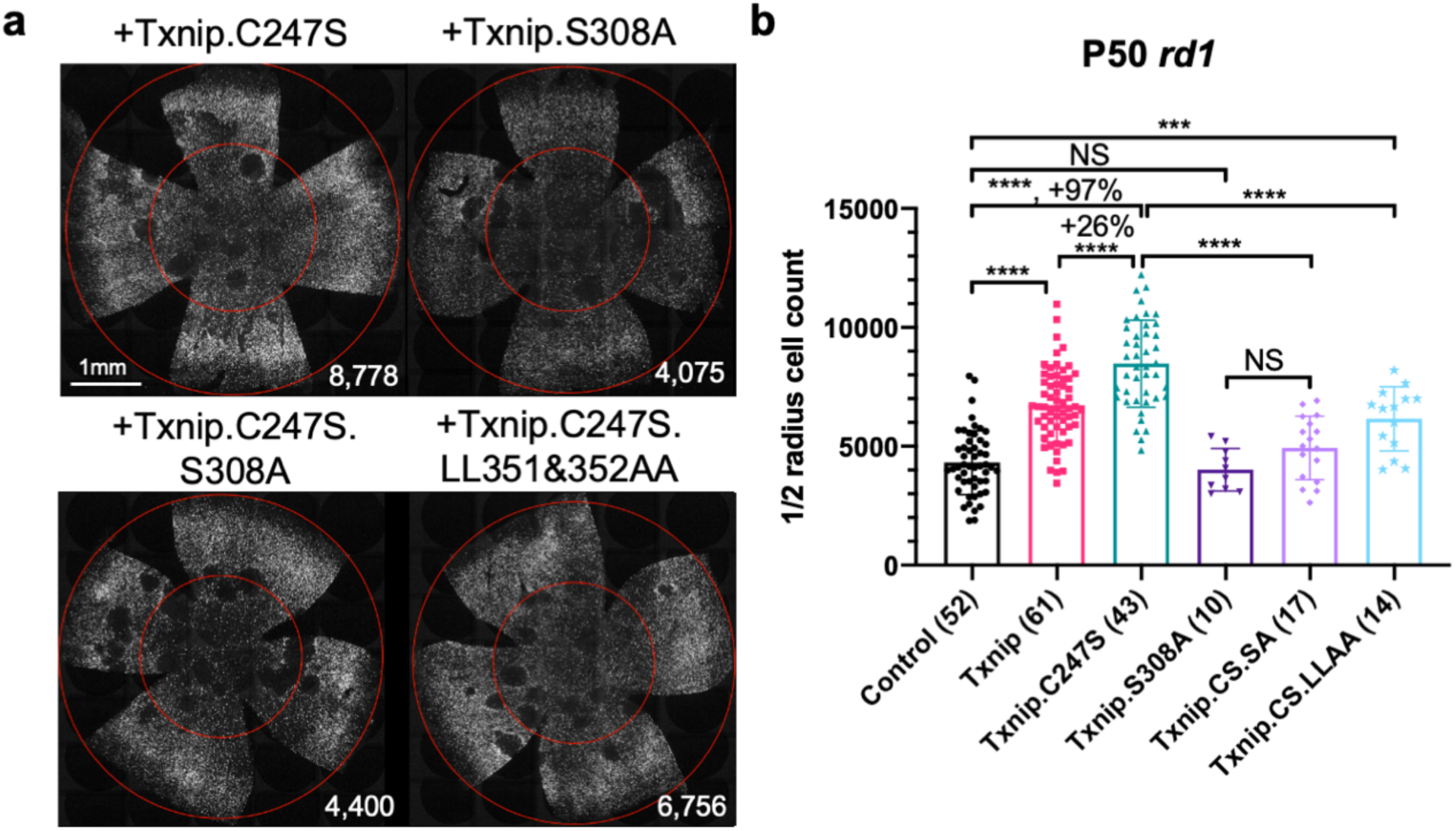
Test of Txnip alleles on cone survival. **a.** Representative P50 *rd1* flat-mounted retinas with H2BGFP (gray) labeled cones treated with one of four different Txnip alleles. **b.** Quantification of H2BGFP-positive cones within the half radius of P50 *rd1* retinas treated with wildtype (wt) Txnip, Txnip alleles, and control. Error bar: standard deviation. Abbreviations: Txnip.CS.SA = Txnip.C247S.S308A; Txnip.CS.LLAA = Txnip.C247S.LL351&352AA. NS: not significant, *p* > 0.05. * *p* < 0.05. ** *p* < 0.01. *** *p* < 0.001. **** *p* < or << 0.0001.

### Txnip requires lactate dehydrogenase b (Ldhb) to prolong cone survival

Human carrying Txnip null mutant presents lactic acidosis (Katsu-Jiménez et al., 2019), suggesting Txnip deficiency compromises lactate catabolism. A recent metabolomic study of muscle using a targeted knock-out of Txnip suggested that Txnip increases the catabolism of non-glucose fuels, such as lactate, ketone bodies and lipids (DeBalsi et al., 2014). This switch in fuel preference was proposed to benefit the mitochondrial tricarboxylic acid cycle (TCA) cycle, leading to a greater production of ATP. As presented earlier, a problem for cones in the RP environment might be a shortage of glucose (Aït-Ali et al., 2015; Punzo et al., 2009; Wang et al., 2016). A benefit of Txnip might then be to enable and/or force cells to switch from a preference for glucose to one or more alternative fuels. To test this hypothesis, we co-injected AAV-Txnip with shRNAs targeting the rate-limiting genes for the catalysis of lactate, ketones or lipids. Ldhb, encoded by *ldhb* gene, is the enzyme that converts lactate to pyruvate to potentially fuel the TCA cycle, and lactate dehydrogenase a (Ldha, encoded by *ldha* gene), converts pyruvate to lactate (Eventoff et al., 1977). We found that Txnip rescue was significantly decreased by any one of three Ldhb shRNAs (siLdhb) or by overexpression of Ldha (Fig. 3a,b and Extended Data Fig. 3). We also tested the rescue effect of Txnip plus an shRNA against Oxct1 (siOxct1), a critical enzyme for ketolysis (Zhang and Xie, 2017), or against Cpt1a (siCpt1a), a component for lipid transporter that is rate limiting for β-oxidation (Shriver and Manchester, 2011). These shRNAs, tested singly or in combination, did not reduce the effectiveness of Txnip rescue (Fig. 3c). Taken together, these data support the use of lactate, but not ketones or lipids, as a critical alternative fuel for cones when Txnip is overexpressed.

**Fig. 3:**
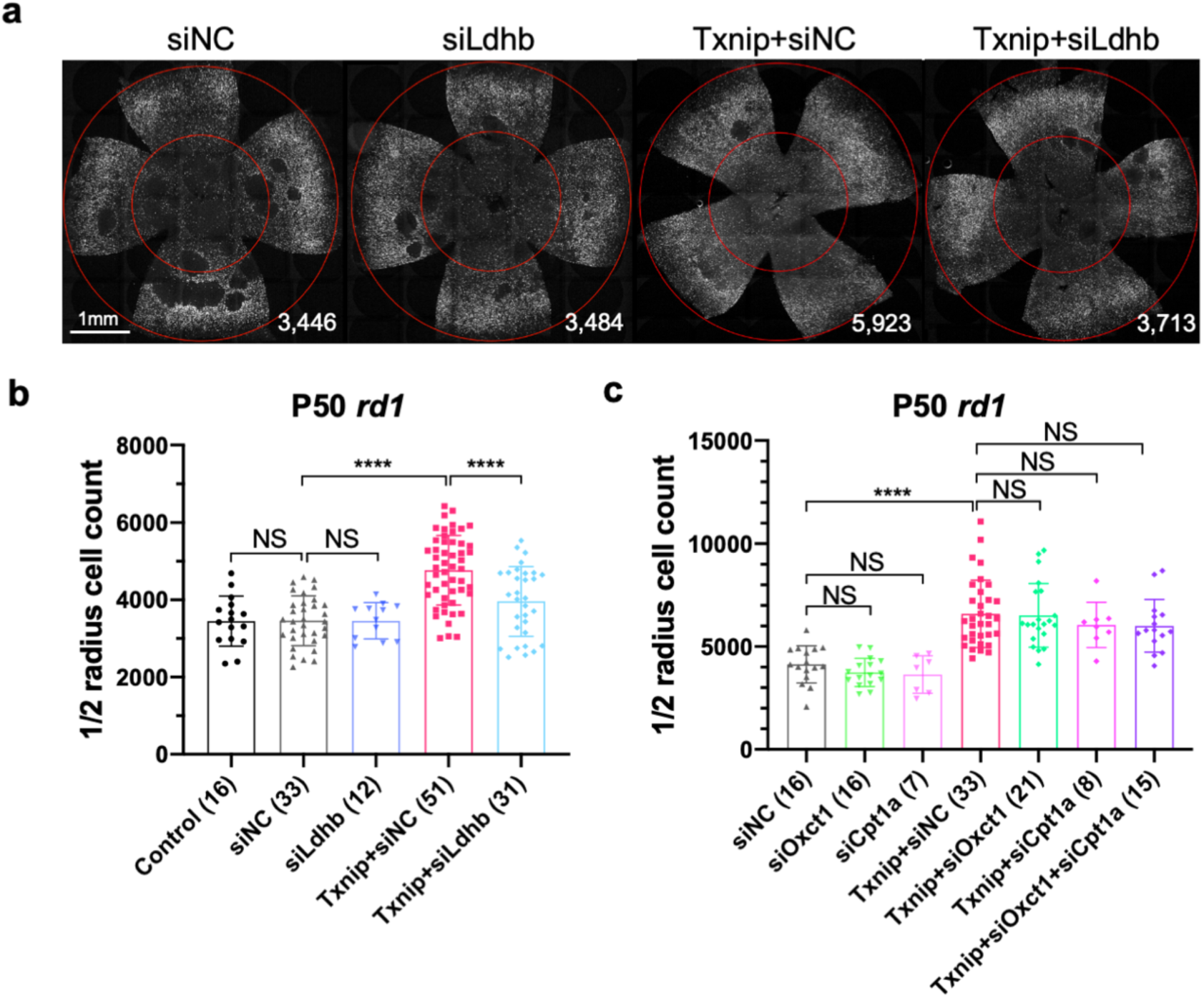
Ldhb is necessary for Txnip-induced rescue of RP cones *in vivo*. **a.** Representative P50 *rd1* flat-mounted retinas with H2BGFP (gray) labeled cones treated with siNC (non-targeting scrambled control shRNA), siLdhb^(#2)^ (Ldhb shRNA), Txnip + siNC, or Txnip + siLdhb^(#2)^. **b.** Quantification of H2BGFP-positive cones within the half radius of P50 *rd1* retinas treated with control, siNC control, Txnip + siLdhb^(#2)^ or siNC control. **c.** Quantification of H2BGFP-positive cones within the half radius of P50 *rd1* retinas treated with Txnip + siOxct1^(#c)^, Txnip + siCpt1a^(#c)^, Txnip + siOxct1^(#c)^ + siCpt1a^(#c)^, or siNC control. Error bar: standard deviation. NS: not significant, *p* > 0.05. ** *p* < 0.01. *** *p* < 0.001. **** *p* < or << 0.0001.

### Txnip improves the ATP:ADP ratio in RP cones in the presence of lactate

If the improved survival of cones following Txnip overexpression is due to improved utilization of non-glucose fuels, cones might show improved mitochondrial metabolism. To begin to examine the metabolism of cones, we first attempted to perform metabolomics of cones with and without Txnip. However, so few cones are present in these retinas that we were unable to achieve meaningful results. An alternative assay was conducted to measure the ratio of ATP to ADP using a genetically-encoded fluorescent sensor (GEFS). AAV was used to deliver PercevalHR, an ATP:ADP GEFS (Tantama et al., 2013), to *rd1* cones with and without AAV-Txnip. The infected P20 *rd1* retinas were explanted and imaged in three different types of media to measure the cone cytosolic ratio of ATP:ADP. Txnip increased the ATP:ADP ratio (i.e. higher F_PercevalHR_^488:405^) of *rd1* cones in lactate-only or pyruvate-only media. Consistent with the role of Txnip in removing Glut1 from the plasma membrane, Txnip treated cones had a lower ATP:ADP ratio (i.e. lower F_PercevalHR_^488:405^) in high glucose medium (Fig. 4a,b). To further probe whether intracellular glucose was reduced after overexpression of Txnip (Wu et al., 2013), a glucose sensor iGlucoSnFR was used (Keller et al., 2019). This sensor showed reduced intracellular glucose in Txnip-treated cones (Extended Data Fig. 4a,b). Because the fluorescence of GEFS may be also subject to environmental pH, we used a pH sensor, pHRed (Tantama et al., 2011), to determine if the changes of PercevalHR and iGlucoseSnFR were due to a change in pH, and it showed no significant pH change (Extended Data Fig. 4c,d). We also found that lactate, but not pyruvate, utilization by Txnip-treated cones was critically dependent upon Ldhb for ATP production, as introduction of siLdhb abrogated the increase in ATP:ADP in Txnip-treated cones (Fig. 4c). Furthermore, in correlation with improved cone survival by Txnip.C247S compared to wt Txnip (Fig. 2b), cones had a higher ATP:ADP ratio in lactate medium when Txnip.C247S was used relative to wt Txnip (Fig. 4e). Similarly, in correlation with no survival benefit when treated with Txnip.S308A (Fig. 2b), there was no difference in the ATP:ADP ratio when Txnip.S308A was used, relative to control, in lactate medium (Fig. 4e).

**Fig. 4:**
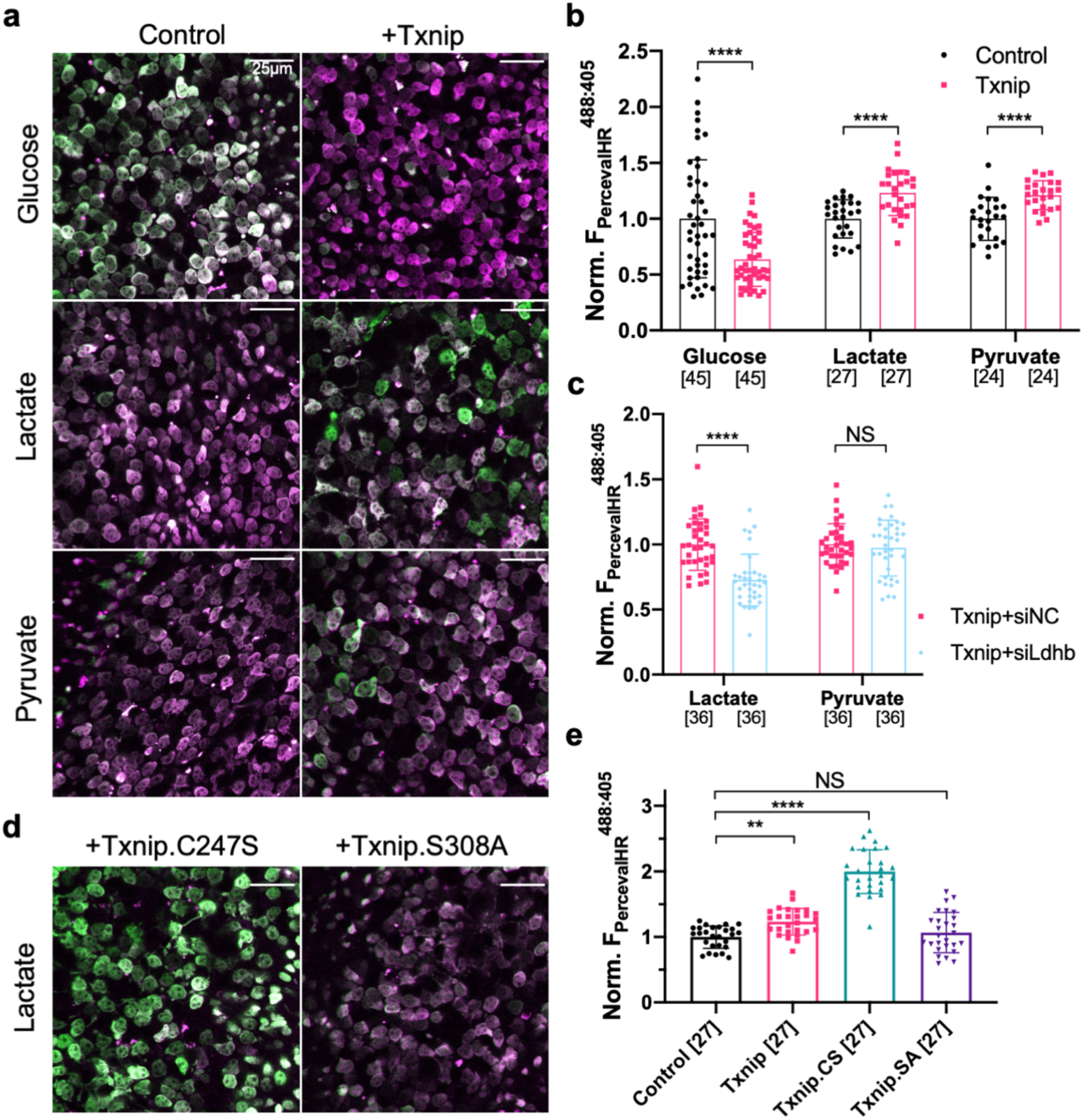
Txnip increases ATP:ADP levels in RP cones in lactate medium. **a.** Representative *ex vivo* live images of PercevalHR labeled cones in P20 *rd1* retinas cultured with high-glucose, lactate-only, or pyruvate-only medium and treated with Txnip or control. (Magenta: fluorescence by 405 nm excitation, indicating low-ATP:ADP. Green: fluorescence by 488 nm excitation, indicating high-ATP:ADP.) **b.** Quantification of normalized PercevalHR fluorescence intensity ratio (F_PercevalHR_^ex488nm : ex405nm^, proportional to ATP:ADP ratio) in cones from P20 *rd1* retinas in different conditions. The number in the bracket “[]” indicates the sample size, i.e. the number of images taken from regions of interest of multiple retinas (≈3 images per retina), in each condition (same applies to all other figures). **c.** Quantification of normalized PercevalHR fluorescence intensity of Txnip + siLdhb^(#2)^ and Txnip + siNC in cones from P20 *rd1* retina in lactate-only or pyruvate-only medium. **d.** Representative *ex vivo* live images of PercevalHR labeled cones in P20 *rd1* retinas cultured in lactate-only medium, following treatment with Txnip.C247S or Txnip.S308A. (Magenta: fluorescence by 405 nm excitation, indicating low-ATP:ADP. Green: fluorescence by 488 nm excitation, indicating high-ATP:ADP.) **e.** Quantification of normalized PercevalHR fluorescence intensity following treatment by Txnip, Txnip alleles, and control cones in *P20 rd1* retinas cultured in lactate-only medium. Abbreviations: Txnip.CS = Txnip.C247S; Txnip.SA = Txnip.S308A. Error bar: standard deviation. NS: not significant, *p* > 0.05. ** *p* < 0.01. *** *p* < 0.001. **** *p* < or << 0.0001.

### Txnip improved RP cone mitochondrial gene expression, size, and function

To further probe the mechanism(s) of Txnip rescue, we first tested if all of the benefits of Txnip were due to Txnip’s effects on Ldhb. Ldhb was thus overexpressed alone or with Txnip. Ldhb alone did not prolong cone survival, nor did it increase the Txnip rescue (Extended Data Fig. 6e). An additional experiment was carried out to investigate if there might be a shortage of the mitochondrial pyruvate carrier, which could limit the uptake of pyruvate into the mitochondria of photoreceptors for ATP synthesis (Grenell et al., 2019). The pyruvate carrier, which is a dimer encoded by *mpc1* and *mpc2* genes, thus was overexpressed, but did not prolong *rd1* cone survival (Extended Data Fig. 6c). To take a less biased approach, the transcriptomic differences between Txnip-treated and control RP cones were characterized. H2BGFP labeled RP cones were isolated by FACS-sorting at an age when cones were beginning to die, and RNA-sequencing was performed (Extended Data Fig. 5a). Data were obtained from two RP strains, *rd1* and *rho*^-/-^. By comparing the differentially expressed genes in common between the two strains, relative to control, seven genes were seen to be upregulated and 17 were downregulated (Supplementary Table 2). Three of the seven upregulated genes were mitochondrial electron transport chain (ETC) genes. The upregulation of these three ETC genes in Txnip-treated *rd1* cones was confirmed by ddPCR (Extended Data Fig. 5b).

The finding of upregulated ETC genes in Txnip-treated cones suggested effects on mitochondria, and thus the morphology of Txnip-treated mitochondria in RP cones was examined by electron microscopy (EM). There was an increase in mitochondrial size by Txnip treatment, with a greater increase in size following treatment with Txnip.C247S (Fig. 5a,b). Mitochondrial membrane potential (ΔΨm) activity, a reflection of mitochondrial ETC function, was also examined using JC-1 dye staining of freshly explanted Txnip-treated P20 *rd1* retinas (Reers et al., 1995). Both Txnip and Txnip.C247S increased the ratio of J-aggregates:JC1-monomers (Fig. 5c,d), indicating an increased ΔΨm and/or a greater number/size of mitochondria with a high ΔΨm following Txnip overexpression. This finding was further investigated *in vivo* using infection by an AAV encoding mitoRFP, which only accumulates in mitochondria with a high ΔΨm (Brodier et al., 2020; Hood et al., 2003). Compared to the control cones without Txnip treatment, the intensity of mitoRFP was higher in P20 *rd1* cones treated with Txnip (Extended Data Fig. 5c,d).

**Fig. 5:**
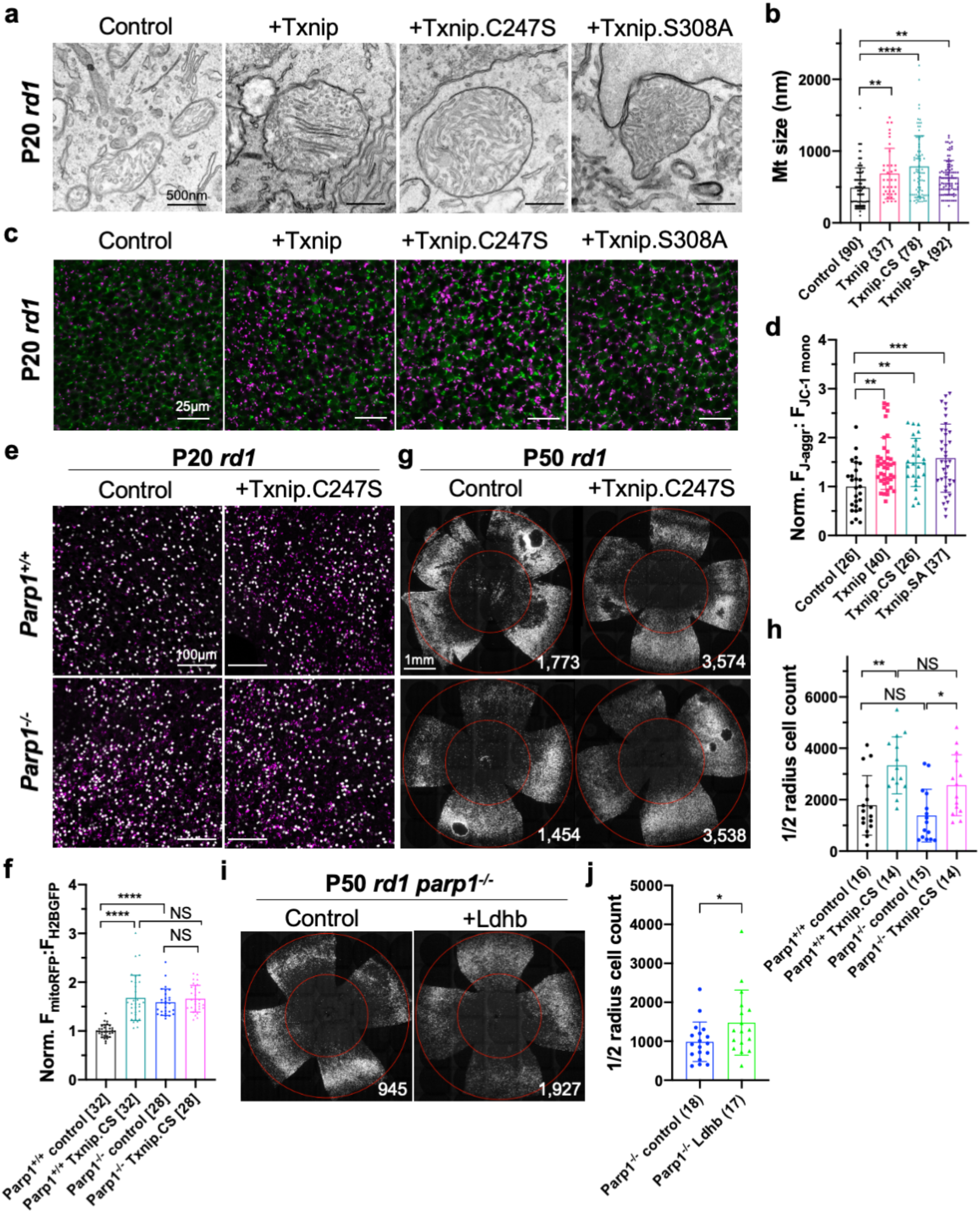
Txnip enhances RP cone mitochondrial size and function. **a.** Representative EM images of RP cones from P20 *rd1* cones treated with Txnip, Txnip.C247S, Txnip.S308A, and control. **b.** Quantification of mitochondrial diameters from control, Txnip, Txnip.C247S and Txnip.S308A treated cones. The number in the curly bracket “{ }” indicates the sample size, i.e. the number of mitochondria from multiple cones of ≥ one retina for each condition (5 retinas for control, 4 for Txnip, 2 for Txnip.C247S, and 1 for Txnip.S308A). **c.** Images of JC-1 dye staining (indicator of ETC function) in live cones of P20 *rd1* central retina at different conditions. (Magenta: J-aggregate, indicating high ETC function. Green: JC-1 monomer, for self-normalization. H2BGFP channel, the tracer of AAV infected area, is not shown.) **d.** Quantification of normalized cone JC-1 dye staining (fluorescence intensity of J-aggregate:JC-1 monomer) from live cones in P20 *rd1* retinas in different conditions (3 - 4 images per retina). **e.** Images of mitoRFP staining (reflecting mitochondrial function) in Txnip.C247S and control cones from fixed P20 *parp1*^+/+^ *rd1* and *parp1*^-/-^ *rd1* retinas near the optic nerve head. (Magenta: mitoRFP. Gray: H2BGFP, for mitoRPF normalization.) **f.** Quantification of normalized mito-RFP:H2BGFP intensity in different conditions of P20 *parp1 rd1* retinas (4 images per retina, near optical nerve head). **g.** Images of P50 *parp1*^+/+^ *rd1* and *parp1*^-/-^ *rd1* retinas with H2BGFP (gray) labeled cones treated with Txnip.C247S or control. *Rd1* cone degeneration seems to be faster after being crossed to *parp1* mice (on 129S background) due to unknown reason(s). Please enlarge the images to appreciate the improved cone counts near the inner circle. **h.** Quantification of H2BGFP-positive cones within the half radius of P50 *parp1*^+/+^ *rd1* and *parp1*^-/-^ *rd1* retinas treated with Txnip.C247S or control. **i.** Images of P50 *parp1*^-/-^ *rd1* retinas with H2BGFP (gray) labeled cones treated with Ldhb or H2BGFP only. **j.** Quantification of H2BGFP-positive cones within the half radius of P50 *parp1*^-/-^ *rd1* retinas treated with Ldhb or H2BGFP only. Error bar: standard deviation. NS: not significant, *p* > 0.05. * *p* < 0.05. ** *p* < 0.01. *** *p* < 0.001. **** *p* < or << 0.0001. Abbreviations: Txnip.SA = Txnip; Txnip.SA = Txnip.S308A.

A previous study identified 15 proteins that interact with Txnip.C247S (Forred et al., 2016). Among these interactors was Parp1, which can negatively affect mitochondria through deleterious effects on the mitochondrial genome (Hocsak et al., 2017; Szczesny et al., 2014), as well as have effects on inflammation and other cellular pathways (Fehr et al., 2020). Due to the similarities between the effects of Txnip addition and of Parp1 inhibition on mitochondria, Parp1 was tested for a potential role in Txnip-mediated rescue. Parp1 expression was first examined by immunohistochemistry and found to be enriched in cone inner segments, which are packed with mitochondria (Hoang et al., 2002), and in cone nuclei (Extended Data Fig. 5g). Interestingly, these are the same locations where a GFP-Txnip fusion protein was found (Extended Data Fig. 1b). To test for a role of Parp1, *parp1*^-/-^ mice were bred to *rd1* mice, and their cone mitochondria were examined by EM and mitoRFP. *Parp1*^-/-^ *rd1* cones possessed larger mitochondria (Extended Data Fig. 5h,i) and higher mitoRFP signals than cones from *parp1*^+/+^ *rd1* controls. Addition of Txnip.C247S to *parp1*^-/-^ *rd1* cones did not alter the mitoRFP signals (Fig. 5e,f). However, when Txnip.C247S was added to *parp1*^-/-^ *rd1* retinas, cone survival was similar to that of Txnip.C247S-treatred *parp1*^+/+^ *rd1* retinas, showing that Txnip-mediated survival does not require Parp1 (Fig. 5g,h).

The discordance between improved mitochondria and cone survival in these experiments suggested that mitochondrial improvement alone is not sufficient to prolong cone survival. This is consistent with the observations from treatment with Txnip.S308A, as well as Txnip + siLdhb, both of which failed to prolong *rd1* cone survival despite improvements in mitochondria (Fig. 2a,b,5a,b,c,d, and Extended Data Fig. 5c,d,e,f). To test if improved cone survival requires both mitochondrial improvement and enhanced lactate catabolism, we delivered Ldhb to *parp1*^-/-^ *rd1* cones. A small but significant improvement in cone survival was observed (Fig. 5i,j).

### Txnip enhances Na^+^/K^+^ pump function and cone opsin expression

The results above suggest that Txnip may prolong RP cone survival by enhancing lactate catabolism via Ldhb, which may lead to greater ATP production by the oxidative phosphorylation (OXPHOS) pathway. Cone photoreceptors are known to require high levels of ATP to maintain their membrane potential, relying primarily upon the Na^+^/K^+^ ATPase pump (Ingram et al., 2020). To investigate whether Txnip affects the function of the Na^+^/K^+^ pump in RP cones, freshly explanted P20 *rd1* retinas were treated with RH421, a fluorescent small-molecule probe for Na^+^/K^+^ pump function (Fedosova et al., 1995). Addition of Txnip improved Na^+^/K^+^ pump function of these cones in lactate medium as reflected by an increase in RH421 fluorescence (Fig. 6a,b), consistent with Txnip enabling greater utilization of lactate. In RP cones, it is also known that protein expression of cone opsin is down-regulated, postulated to be due to insufficient energy supply (Punzo et al., 2009). Compared to control, greater anti-opsin staining was observed in Txnip-treated *rd1* cones at P50 (Fig. 6c), further supporting the idea that Txnip improves the energy supply to RP cones.

**Fig. 6:**
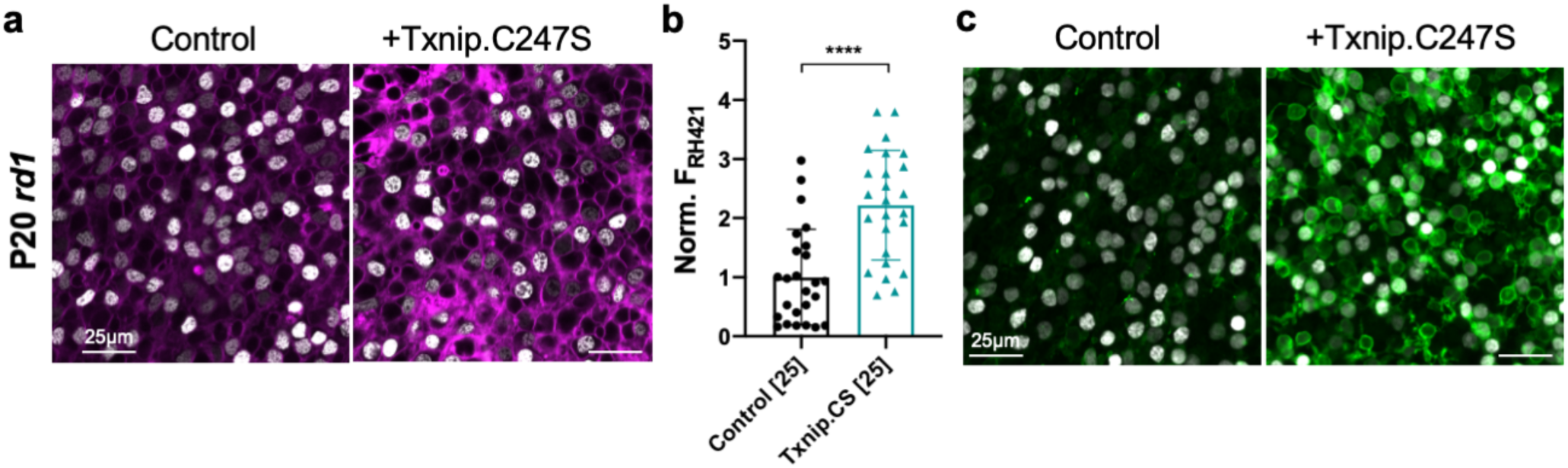
Txnip enhances Na^+^/K^+^ ATPase pump function and cone opsin expression in RP cones. **a.** Images of live *ex vivo* RH421 stained cones in P20 *rd1* retinas treated with Txnip.C247S or control and cultured in lactate-only medium. (Magenta: RH421 fluorescence, proportional to Na^+^/K^+^ ATPase function. Gray: H2BGFP, tracer of infection). **b.** Quantification of normalized RH421 fluorescence intensity from Txnip.C247S treated cones relative to control in P20 *rd1* retinas cultured in lactate-only medium (5 images per retina). Abbreviation: Txnip.CS = Txnip.C247S. **c.** IHC with anti-s-opsin plus anti-m-opsin antibodies near the half radius of P50 *rd1* retinas treated with Txnip.C247S or control. (Green: cone-opsins. Gray: H2BGFP, tracer of infection). Error bar: standard deviation. **** *p* < or << 0.0001.

### Dominant-negative HIF1α improves RP cone survival

If improved lactate catabolism and OXPHOS are at least part of the mechanism of Txnip rescue, RP cone survival might be promoted by other molecules serving similar functions. HIF1α can upregulate the transcription of glycolytic genes (Majmundar et al., 2010). Increased glycolytic enzyme levels might push RP cones to rely on glucose, rather than lactate, to their detriment if glucose is limited. To investigate whether HIF1α might play a role in cone survival, a wt and a dominant-negative HIF1α (dnHIF1α) allele (Jiang et al., 1996) were delivered to *rd1* retinas using AAV. A target gene of HIF1α, *vegf*, which might improve blood flow and thus nutrient delivery, also was tested. The dnHIF1α increased *rd1* cone survival, while wt HIF1α and Vegf each decreased cone survival (Fig. 7a,b, Extended Data Fig. 6d,e).

**Fig. 7:**
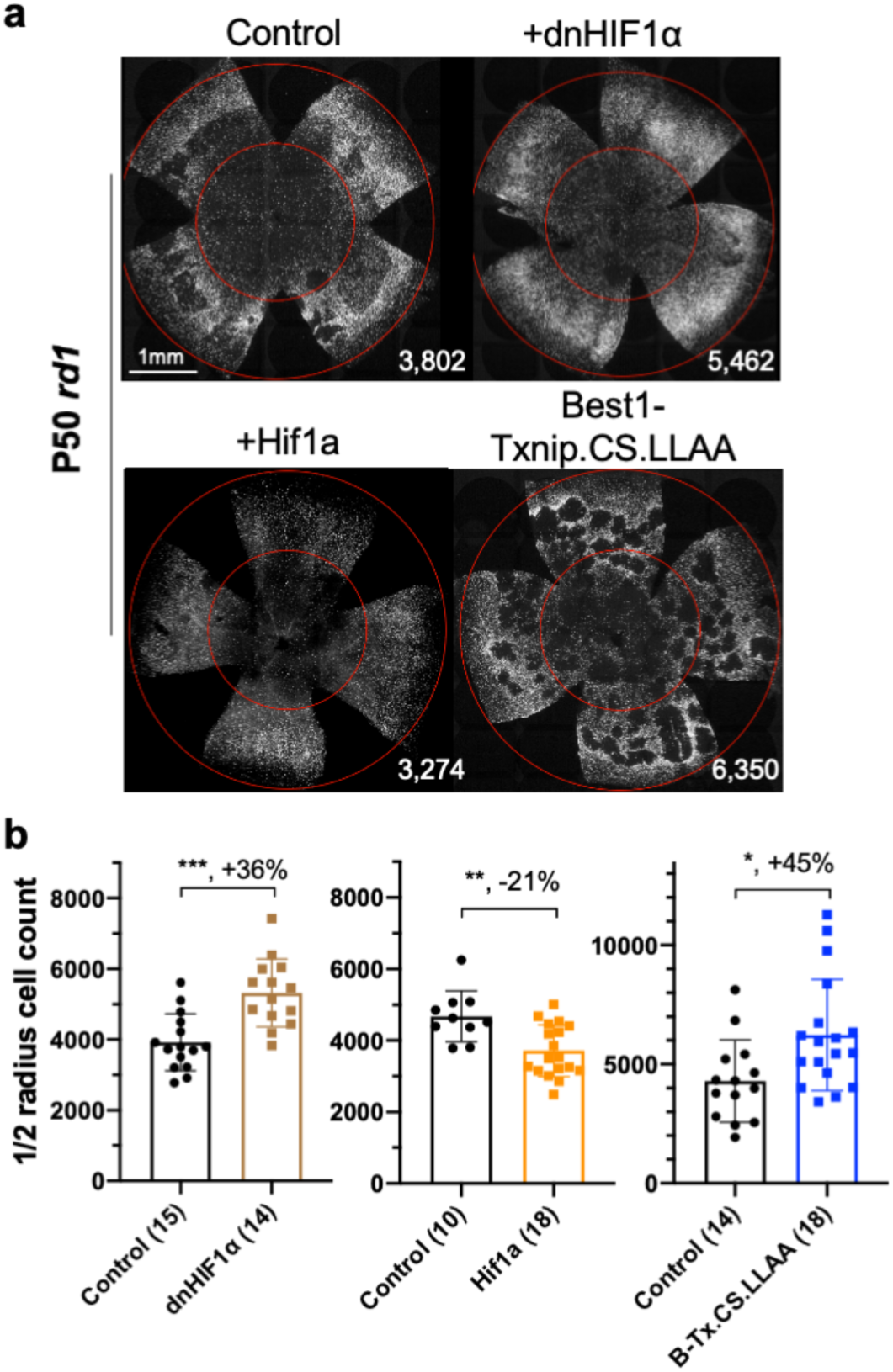
Dominant negative HIF1α and Best1-Txnip.C247S.LL351&352AA enhance RP cone survival. **a.** Images of P50 *rd1* retinas with H2BGFP (gray) labeled cones treated with dnHIF1α, Hif1a, Best1-Txnip.C247S.LL351&352AA (Txnip.CS.LLAA, driven by an RPE-specific promoter) or control. Note that Best1-Txnip.C247S.LL351&352AA amplified the FVB-specific retinal craters, while dnHIF1α decreased them. **b.** Quantification of H2BGFP-positive cones within the half radius of P50 *rd1* retinas treated with dnHIF1α, Hif1a, Best1-Txnip.C247S.LL351&352AA or control. Abbreviation: B-Tx.CS.LLAA = Best1-Txnip.C247S.LL351&352AA. Error bar: standard deviation. * *p* < 0.05. ** *p* < 0.01. *** *p* < 0.001.

### Txnip effects on Glut1 levels in the RPE and cone survival

Several lines of evidence support the hypothesis that RP cones do not have sufficient glucose to satisfy their needs via glycolysis (Chinchore et al., 2019; Kanow et al., 2017; Punzo et al., 2012, 2009; Wang et al., 2016). To determine if retention of glucose by the RPE might underlie a glucose shortage for cones (Kanow et al., 2017; Wang et al., 2016), we attempted to reprogram RPE metabolism to a more “OXPHOS” and less “glycolytic” status by overexpressing Txnip or dnHIF1α with an RPE-specific promoter, the Best1 promoter (Esumi et al., 2009). The goal was to increase lactate consumption in the RPE, thus freeing up more glucose for delivery to cones. However, no RP cone rescue was observed (Extended Data Fig. 6b), possibly due to a clearance of Glut1 from the surface of cells, which would create a glucose shortage for both the RPE and the cones (Swarup et al., 2019) (Extended Data Fig. 6a). To examine the level of Glut1 in the RPE following introduction of wt Txnip, or Txnip.C247S.LL351&352AA, which should prevent efficient removal of Glut1, immunohistochemistry for Glut1 was carried out. This assay showed that AAV-Best1-Txnip.LL351&352AA did result in less clearance of Glut1 from the surface of the RPE (Extended Data Fig. 6a) relative to wt Txnip. Txnip.C247S.LL351&352AA was then tested for *rd1* cone rescue, where it was found to improve cone survival (Fig. 7a,b), in keeping with the model that the RPE retains glucose to the detriment of cones in RP.

### Combination of Txnip.C247S with other rescue genes provides an additive effect

Finally, as our goal is to provide effective, generic gene therapy for RP, and potentially other diseases that affect photoreceptor survival, we used combinations of AAVs that encode genes that we have previously shown prolong RP cone survival and vision. The combination of Txnip.C247S expression in cones, with expression of Nrf2, a gene with anti-oxidative damage and anti-inflammatory activity, in the RPE, provided an additive effect on cone survival relative to either gene alone (Fig. 8a,b). This combination also preserved the RP cone outer segments, which is the structure packed with opsin for photon detection, and reduced the mislocalization of opsin to the plasma membrane (Fig. 8c). An interesting phenotype that is especially prominent in the FVB *rd1* strain is that of “craters” in the photoreceptor layer. These are areas of circumscribed cone death that are obvious when the retina is viewed as a flat-mount. AAV-Best1-Nrf2 alone suppressed the formation of these craters (Wu et al., n.d.), while AAV-RedO-Txnip did not, despite the fact that AAV-RedO-Txnip.C247S provides the most robust RP cone rescue that we have seen (Fig. 2a,6f,6h). An additional combination that was tested was AAV-RedO-Txnip.C247S with AAV-RedO-Tgfb1, an anti-inflammatory gene (Wang et al., 2020). This combination did not improve cone survival beyond that of Txnip alone, but almost completely eliminated the craters (Fig. 8d,e). In addition, we tried other genes in combination with wt Txnip, but did not observe any obvious improvement over Txnip alone (Extended Data Fig. 6e).

**Fig. 8:**
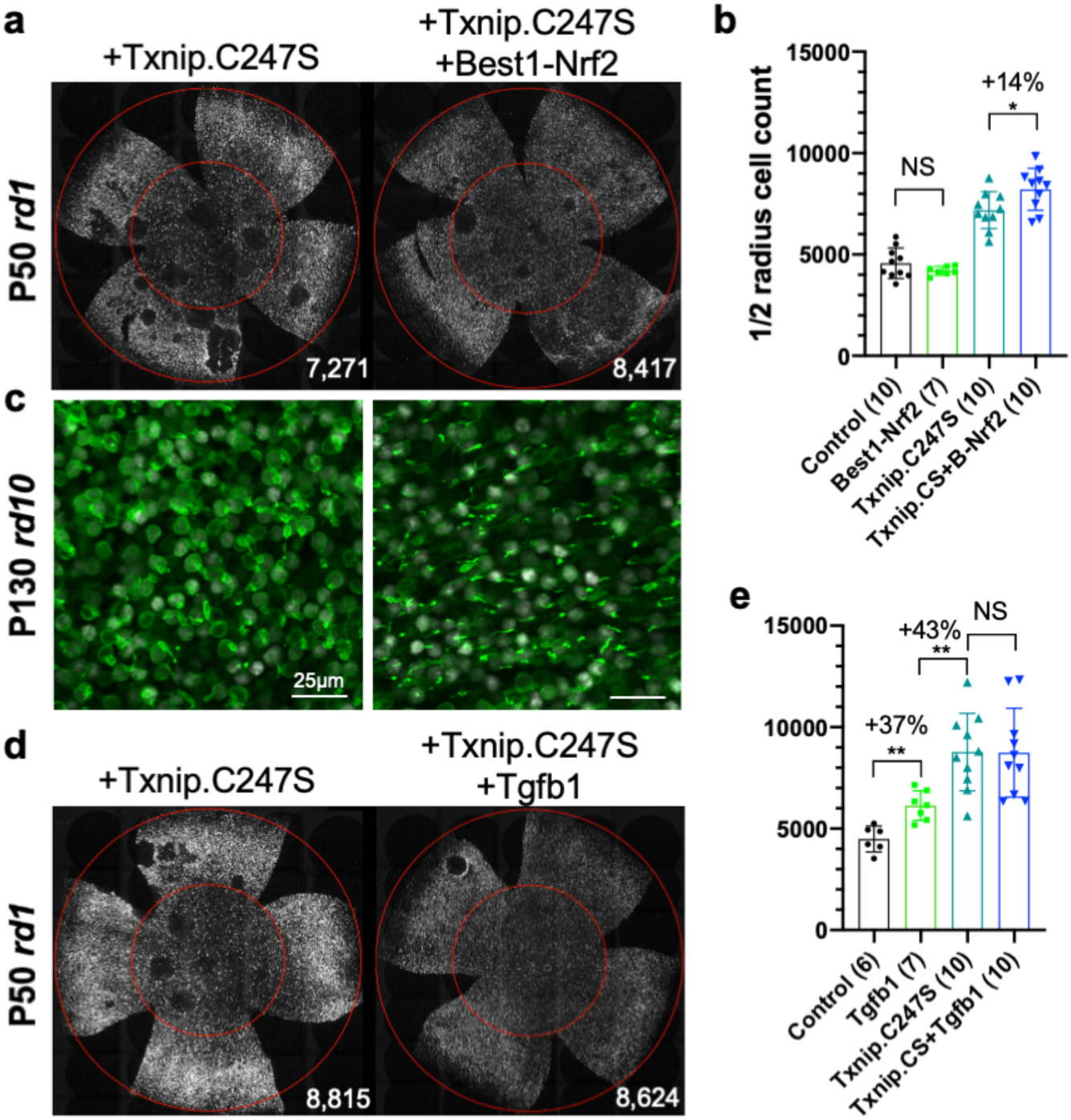
Combination of Txnip.C247S with Best1-Nrf2 or Tgfb1 provides an additive effect. **a.** Images of P50 rd1 retinas with H2BGFP (gray) labeled cones treated with Txnip.C247S or Txnip.C247S + Best1-Nrf2. **b.** Quantification of H2BGFP-positive cones within the half radius of P50 *rd1* retinas treated with Txnip.C247S or Txnip.C247S + Best1-Nrf2. Abbreviation: B-Nrf2 = Best1-Nrf2. **c.** IHC with anti-s-opsin plus anti-m-opsin antibodies near the half radius of P130 *rd10* retinas treated with Txnip.C247S (left panel) or Txnip.C247S + Best1-Nrf2 (right panel). (Green: cone-opsins. Gray: H2BGFP, tracer of infection). **d.** Images of P50 rd1 retinas with H2BGFP (gray) labeled cones treated with Txnip.C247S or Txnip.C247S + Tgfb1. **e.** Quantification of H2BGFP-labeled cones within the half radius of P50 *rd1* retinas treated with control, Tgfb1, Txnip.C247S, or Txnip.C247S + Tgfb1. Error bar: standard deviation. NS: not significant, *p* > 0.05. * *p* < 0.05. ** *p* < 0.01.

## Discussion

Photoreceptors have been characterized as being highly glycolytic, even under aerobic conditions, as originally described by Otto Warburg (Warburg, 1925). Glucose appears to be supplied primarily from the circulation, via the RPE, which has a high level of Glut1 (Gospe et al., 2010). Photoreceptors, at least rods, carry out glycolysis to support anabolism, to replace their outer segments (Chinchore et al., 2017), and contribute ATP, to run their ion pumps (Okawa et al., 2008). If glucose becomes limited, as has been proposed to occur in RP, cones may have insufficient fuel for their needs. To explore whether we could develop a therapy to address some of these metabolic shortcomings in RP, we delivered many different types of genes that might alter metabolic programming. From these, Txnip had the strongest benefit on cone survival and vision (Fig. 1 and Extended Data Fig. 1). This was surprising as Txnip has been shown to inhibit glucose uptake, by binding to and aiding in the removal of Glut1 from plasma membrane, and it inhibits the anti-oxidation proteins, the thioredoxins, again by direct binding. The results with Txnip in its wt form, and from the study of several mutant alleles, provide some insight into how it might benefit cones. The Txnip.C247S allele prevents binding to thioredoxins, and gave enhanced cone survival relative to wt Txnip (Fig. 2 and Extended Data Fig. 2). We speculate that, by being free of this interaction, the C247S mutant protein may be more available for other Txnip-mediated activities. In addition, thioredoxin may be made more available for its role in fighting oxidative damage.

The mechanisms by which Txnip might benefit cones are not fully known, but a study of Txnip’s function in skeletal muscle suggested that it plays a role in fuel selection (DeBalsi et al., 2014). If glucose is limited in RP, then cones may need to switch from a reliance on glucose and glycolysis to an alternative fuel(s), such as ketones, fatty acids, amino acids, or lactate. Cones express *oxct1* mRNA (Shekhar et al., 2016), which encodes a critical enzyme for ketone catabolism, suggesting cones are capable of ketolysis. In addition, a previous study showed that lipids might be an alternative energy source for cones by β-oxidation (Joyal et al., 2016). It is likely that cones can use these alternative fuels to meet their intense energy demands (Ingram et al., 2020) (Fig. 6,7). However, the Txnip rescue did not depend on ketolysis or β-oxidation (Fig. 3). Due to the diversity of amino acid catabolic pathways, we did not study whether these pathways were required for Txnip’s rescue effect. However, we did discover that Ldhb, which converts lactate to pyruvate, was required. This is an interesting switch, as photoreceptors normally have high levels of Ldha, and produce lactate (Chinchore et al., 2017). An important factor in the reliance on Ldhb could be the availability of lactate, which is highly available from serum (Hui et al., 2017). Lactate could be transported via the RPE and/or Müller glia, and/or the internal retinal vasculature which comes in closer proximity to cones after rod death. Ketones are usually only available during fasting, and lipids are hydrophobic molecules which are slow to be transported across the plasma membranes. Moreover, lipids are required to rebuild the membrane-rich outer segments, and thus might be somewhat limited. Ldhb is not sufficient, however, to delay RP cone degeneration, as its overexpression did not promote RP cone survival.

Txnip-treated RP cones also had larger mitochondria with a greater membrane potential, and likely were able to use the pyruvate produced by Ldhb for greater ATP production via OXPHOS. Indeed, Txnip-treated cones had an enhanced ATP:ADP ratio (Fig. 4). However, healthier mitochondria were not sufficient to prolong RP cone survival. Txnip.S308A led to larger mitochondria than control mitochondria, brighter JC-1 staining and mitoRFP signals, which are indicators of better mitochondrial health, but this allele did not induce greater cone survival (Fig. 5 and Extended Data Fig. 5). Moreover, as Txnip has been shown to interact with Parp1, which can negatively affect mitochondria, we investigated if Parp1 knock-out mice might have cones that survive longer in RP. Indeed, the Parp1 knock-out mitochondria appeared to be healthier, but Parp1 knock-out retinas did not have better RP cone survival than Parp1-wt *rd1* retinas. In addition, cone rescue by Txnip was not changed in the Parp1 knock-out retinas.

The well-described effects of Txnip on the removal of Glut1 from the cell membrane might seem at odds with the promotion of cone survival. It could be that removal of Glut1 from the plasma membrane of cones forces the cones to choose an alternative fuel, such as lactate, and perhaps others too. Interestingly, as Glut1 knock-down was not sufficient for cone survival, Txnip must not only lead to a reduction in membrane localized Glut1, but also potentiate a fuel switch, via an unknown mechanism(s) that at least involves an increase of Ldhb activity. A reduction in glycolysis might also lead to a fuel switch. Introduction of dnHIF1α, which should reduce expression of glycolytic enzymes, also benefitted cones, while introduction of wt HIF1α did not (Fig. 7). HIF1α has many target genes, and may alter pathways in addition to that of glycolysis, thus also potentiating a fuel switch once glycolysis is down regulated. An additional finding supporting the notion that the level of glycolysis is important for cone survival was the observation that AAV-Pfkm plus AAV-Hk1 led to a reduction in cone survival (Extended Data Fig. 1e). Phosphorylation of glucose by Hk1 followed by phosphorylation of fructose-6-phosphate by the Pfkm complex commits glucose to glycolysis at the cost of ATP. These AAVs may have promoted the flux of glucose through glycolysis, which may have inhibited a fuel switch, and/or depleted the ATP pool, e.g. if downstream glycolytic intermediates were used for anabolic needs so that ATP production by glycolysis did not occur.

The observations described above suggest that at least two different pathways are required for the promotion of cone survival by Txnip (Extended Data Fig. 8). One pathway requires lactate utilization via Ldhb, but as Ldhb was not sufficient, another pathway is also required. As greater mitochondrial health was observed following Txnip treatment, a second pathway may include the effects on mitochondria. This notion is supported by the observation that the addition of Ldhb to *parp1^-/-^ rd1* cones, which have healthier mitochondria, led to improved cone survival (Fig. 5). Txnip alone may be able to promote cone health by impacting both lactate catabolism and mitochondrial health. There may be additional pathways required as well.

The effects of Txnip alleles expressed only in the RPE provide some support for the hypothesis that the RPE transports glucose to cones for their use, while primarily using lactate for its own needs (Kanow et al., 2017; Swarup et al., 2019). Lactate is normally produced at high levels by photoreceptors in the healthy retina. When rods, which are 97% of the photoreceptors, die, lactate production goes down dramatically. The RPE might then need to retain glucose for its own needs. Introduction of an allele of Txnip, C247S.LL351&352AA, to the RPE provided a rescue effect for cones, while introduction of the wt allele of Txnip to the RPE did not. The LL351&352AA mutations lead to a loss of efficiency of the removal of Glut1 from the plasma membrane, while the C247S mutation might create an even less glycolytic RPE. The combination of these mutations might then allow more glucose to flow to cones. The untreated RP cones seem to be able to use glucose at a high concentration for ATP production, at least in freshly explanted retinas (Fig. 4a). These findings are also consistent with the reported mechanism for cone survival promoted by RdCVF, a factor that is proposed to improve glucose uptake by RP cones, which might be important if glucose is present in low concentration due to retention by the RPE (Aït-Ali et al., 2015; Byrne et al., 2015).

As cones face multiple challenges in the degenerating RP retina, we tested Txnip in combination with genes that we have found to promote cone survival via other mechanisms. The combination of Txnip with vectors fighting oxidative stress (AAV-Best1-Nrf2) or inflammation (AAV-RedO-Tgfb1) supported greater cone survival than any of these treatments alone. These combinations utilize cell type-specific promoters, reducing the chances of side effects from global expression of these genes. Of note, the Nrf2 expression was limited to the RPE, yet was additive for cone survival. This finding is in keeping with the interdependence of photoreceptors and the RPE, which is undoubtably important not only in a healthy retina, but in disease as well.

## Methods

### Animals

*rd1* mice were the albino FVB strain, which carries the *Pde6b^rd1^* allele (MGI: 1856373). BALB/c, CD1, and FVB mice were purchased from Charles River Laboratories. Due to availability, part of the FVB mice were purchased from Taconic, and we did not notice any difference between the two sources in terms of cone degeneration rate. C57BL/6J, *rd10*, and *parp1*^-/-^ mice were purchased from The Jackson Laboratories and bred in house. We crossed the *parp1*^-/-^ mice with FVB mice to generate homozygous *parp1*^-/-^ *rd1* and *parp1*^+/+^ *rd1* mice. Genotyping of these mice was done by Transnetyx (Cordova, TN). The *rho*^-/-^ mice were provided from by Lem (Tufts University, MA) (Lem et al., 1999).

### AAV vector design, authentication, and preparation

Detailed information of all AAVs used in this study is listed in Supplementary Table 1, along with the authentication information. cDNAs of mouse *txnip, hif1a, hk2, ldha, ldhb, slc2a1, bsg1, cpt1a, oxct1, mpc1* and *mpc2*, and human *nrf2,* were purchased from GeneCopeia (Rockville, MD). Mouse *vegf164* cDNA (Robinson and Stringer, 2001) was synthesized by Integrated DNA Technologies (Coralville, Iowa). We obtained the following plasmids as gifts from various depositors through Addgene (Watertown, MA): *hk1, pfkm* and *pkm2* (William Hahn & David Root; #23730, #23728, #23757), *pkm1* (Lewis Cantley & Matthew Vander Heiden; #44241), H2B-GFP (Geoff Wahl; #11680), mitoRFP (i.e. DsRed2-mito, Michael Davidson; #55838), GFP-Txnip (Clark Distelhorst; #18758), W3SL (Bong-Kiun Kaang; #61463), 3xFLAG (Thorsten Mascher, #55180), PercevalHR and pHRed (Gary Yellen; #490820, #31473). The cDNA of mouse RdCVF was a gift from Leah Byrne and John Flannery (UC Berkeley, CA). iGlucoSnFR was provided under a Material Transfer Agreement by Jacob Keller and Loren Looger (Janelia Research Campus, VA). RedO promoter was provided as a gift, and SynPVI and SynP136 promoters were provided under a Material Transfer Agreement, from Botond Roska (IOB, Switzerland). The Best1 promoter was synthesized by lab member, Wenjun Xiong, using Integrated DNA Technologies based on literature (Esumi et al., 2009). Mutated Txnip, dominant-negative HIF1α (Jiang et al., 1996) and RO1.7 promoter (Ye et al., 2016) were created from the corresponding wildtype plasmids in house using Gibson assembly.

All of the new constructs in this study were cloned using Gibson assembly. For example, AAV-RedO-Txnip was cloned by replacing the EGFP sequence of AAV-RedO-EGFP at the NotI/HindIII sites, with the Txnip sequence, which was PCR-amplified from the cDNA vector adding two 20-bp overlapping sequences at the 5’- and 3’-ends. All of the AAV plasmids were amplified using Stbl3 *E. coli* (Thermo Fisher Scientific). The sequences of all AAV plasmids were verified with directed sequencing and restriction enzyme digestion. The key plasmids were triple-verified with Next-Generation complete plasmid sequencing (MGH CCIB DNA Core), which is able to capture the full sequence of the ITR regions. The genome sequence of critical AAVs (i.e. AAV8-RedO-Txnip.C247S and AAV8-RedO-Txnip.S308A) were quadruple-verified with PCR and directed sequencing.

All of the vectors were packaged in recombinant AAV8 capsids using HEK293T cells and purified with iodixanol gradient as previously described (Grieger et al., 2006; Xiong et al., 2015). The titer of each AAV batch was determined using protein gels, comparing viral band intensities with a previously established AAV standard. The concentration of our AAV production usually ranged from 2 x 10^12^ to 3 x 10^13^ gc/mL. Multiple batches of key AAV vectors (e.g. 4 batches of AAV8-RedO-Txnip, and 3 batches of AAV-RedO-siLdhb^(#2)^) were made and tested *in vivo* to avoid any unknown batch effects.

### shRNA

The shRNA plasmids of Ldhb, Slc2a1, Oxct1 and Cpt1a were purchased from GeneCopeia, and they were provided as three or four distinct sequences for each gene, driven by the H1 or U6 promoter. The knockdown efficiency of these candidate shRNA sequences was tested by co-transfecting with CAG-TargetGene-IRES-d2GFP vector in HEK293T cells as previously described (Matsuda and Cepko, 2007; Wang et al., 2014). The GFP fluorescence intensity served as a fast and direct read out of the knockdown efficiency of these shRNAs. Using this method, we selected the following sense strand sequences to knockdown the targeted genes (Extended Data Fig. 8-11): siLdhb^(#2)^ 5’-CCATCATCGTGGTTTCCAACC-3’; siLdhb^(#1)^ 5’-GCAGAGAAATGTCAACGTGTT-3’; siLdhb^(#3)^ 5’-GCCGATAAAGATTACTCTGTG-3’; siSlc2a1^(#a)^ 5’-GGTTATTGAGGAGTTCTACAA-3’; siOxct1^(#c)^ 5’-GGAAACAGTTACTGTTCTCCC-3’; siCpt1a^(#c)^ 5’-GCATAAACGCAGAGCATTCCT-3’; and siNC (non-targeting scrambled control sequence) 5’-GCTTCGCGCCGTAGTCTTA-3’. We cloned the entire hairpin sequence (including a 6-bp 5’-end lead sequence 5’-gatccg-3’, a 7-bp loop sequence 5’-TCAAGAG-3’ between sense and antisense strands, and a > 7-bp 3’end sequence 5’-ttttttg-3’) and packaged them into AAV8-RedO-shRNA using Gibson assembly as described above. To maximize the knockdown efficacy using a Pol II promoter in AAV (Giering et al., 2008), no extra base pair was kept between the RedO promoter and the 5’-end lead sequence of shRNAs. Due to the lack of an adequate Ldhb antibody, we confirmed the *in vivo* Ldhb knockdown efficiency of all three AAV8-RedO-siLdhb vectors by co-injection with an AAV8-Ldhb-3xFLAG vector into wildtype mouse eyes and detection for FLAG immunofluorescence as described in the Histology section below (Extended Data Fig. 3a).

### Subretinal injection

On the day of birth (P0), ≈1 x 10^9^ vg/eye of AAV was injected into the eyes of pups as previously described(Matsuda and Cepko, 2007; Xiong et al., 2015). For all experiments in which cones were quantified, and to provide a means to trace infection (e.g. for immunohistochemistry), 2.5 x 10^8^ vg/eye of AAV8-RedO-H2BGFP was co-injected with other AAVs, or alone as a control. For all other experiments, such as FACS sorting and *ex vivo* live-imaging, 1 x 10^9^ vg/eye of AAV8-SynP136-H2BGFP was co-injected.

### Photopic visual acuity measured for optomotor response

The photopic optomotor response of mice were measured using the OptoMotry System (CerebralMechanics) at a background light of ≈70 cd/m^2^ as previously described (Xiong et al., 2019). The contrast of the grates was set to be 100%, and temporal frequency was 1.5 Hz. The threshold of mouse visual acuity (i.e. maximal spatial frequency) was tested by an examiner without knowledge of the injected or the treatment group. During each test, the direction of movement of the grates (i.e. clockwise or counterclockwise) was randomized, and the spatial frequency of each testing episode was determined by the software. Without knowing the spatial frequency of the moving grates, the examiner reported either “yes” or “no” to the system until the threshold of acuity was determined by the software.

### Histology

Mice were euthanized with CO_2_ and cervical dislocation, and the eye was enucleated. For flat-mounts, retinas were separated from the rest of the eye using a dissecting microscope and were fixed in 4% paraformaldehyde solution for 30 minutes. The retinas were then flat mounted on a glass slide and coverslip. For H2BGFP labeled cone imaging, we used a Keyence microscope with a 10x objective (Plan Apo Lamda 10x/0.45 Air DIC N1) and GFP filter box (OP66836).

For cone opsin antibody staining in whole-mount retinas, after fixation, retinas were blocked for 1 hour in PBS with 5% normal donkey serum and 0.3% Triton X-100 at room temperature. After blocking, retinas were incubated with a mixture of 1:200 anti-s-opsin antibody (AB5407, EMD Millipore) and 1:600 anti-m-opsin antibody (AB5405, EMD Millipore) in the same blocking solution overnight at 4 °C, followed by secondary donkey-anti-rabbit antibody staining (1:1000, Alexa Fluor 594) at room temperature for 2 hours, then flat-mounted on a glass slide and coverslip.

For frozen sections, whole eyes were fixed in 4% paraformaldehyde solution for 2 hours at room temperature, followed by removing the cornea, lens and iris. Then the eye cups went through 15% and 30% sucrose gradient to dehydrate at room temperature, followed by overnight incubation in 1:1 30% sucrose and Tissue-Tek® O.C.T. solution at 4 °C. Eye cups were embedded in a plastic mold, frozen in a −80 °C freezer, and cut into 20 μm or 12 μm thin radial cross-sections which were placed on glass slides. Antibody staining was done similarly to whole-mounts as described above and previously (Wang et al., 2014). PBS with 0.1% Triton X-100, 5% normal donkey serum and 1% bovine serum albumin (BSA) was used as the blocking solution, except for FLAG detection (10% donkey serum and 3% BSA). Glut1 (encoded by *slc2a1* gene) antibody (GT11-A, Alpha Diagnostics) was used at 1:300 dilution, Parp1 antibody (ab227244, Abcam) was used at 1:300 dilution, GFP antibody (ab13970, Abcam) was used at 1:1000 dilution to detect GFP-Txnip, and FLAG antibody (ab1257, Abcam) was used at 1:2000 based on a previous study(Ferrando et al., 2015). If applicable, 1:1000 PNA (CY5 or Rhodamine labeled) for cone extracellular matrix labeling, and 1:1000 DAPI were used to co-stain with secondary antibodies. Stained sections were imaged with a confocal microscope (LSM710, Zeiss) using 20x or 63x objectives (Plan Apo 20x/0.8 Air DIC II, or Plan Apo 63X/1.4 Oil DIC III).

### Automated RP cone counting

The cone-H2BGFP images of whole flat-mounted retinas were first analyzed in ImageJ to acquire the diameter and the center parameters of the sample. We used a custom MATLAB script to automatically count the number of H2BGFP-positive cones in the central half of the retina, since RP cones degenerate faster in the central than the peripheral retina. The algorithm was based on a Gaussian model to identify the centers of labeled cells, and published recently (Wu et al., n.d.). The threshold of peak intensity and the variance of distribution were initially determined using visual inspection, and a comparison to the number of manually counted cones from 6 retinas. The threshold of intensity and variance thus determined were then set at fixed values for all the experiments that used cone quantification. The background intensity did not interfere with the accurate counting on the raw images by this MATLAB script, despite the representative images at low-magnification might look differently.

### Live-imaging of cones on *ex vivo* retinal explants

For JC-1 mitochondrial dye staining, the retina was quickly dissected in a solution of 50% Ham’s F-12 Nutrient Mix (11765054, Thermo Fisher Scientific) and 50% Dulbecco’s Modified Eagle Medium (DMEM; 11995065, Thermo Fisher Scientific) at room temperature. They were then incubated in a culture solution containing 50% Fluorobrite DMEM (A1896701, Thermo Fisher Scientific), 25% heat inactivated horse serum (26050088, Thermo Fisher Scientific) and 25% Hanks’ Balanced Salt Solution (HBSS; 14065056, Thermo Fisher Scientific) with 2 μM JC-1 dye (M34152, Thermo Fisher Scientific) at 37 °C in a 5% CO_2_ incubator for 20 minutes. The retinas were washed in 37 °C culture medium without JC-1 for three times, transferred in a glass-bottom culture dish (MatTek P50G-1.5-30-F) with culture medium, and imaged using a confocal microscope (LSM710 Zeiss), which was equipped with a chamber pre-heated to 37 °C with pre-filled 5% CO_2_. Right before imaging, a cover slip (VWR 89015-725) was gently applied to flatten the retina. Regions of interest (with H2BGFP as an indicator of successful AAV infection and to set the correct focal plane on the cone layer) were selected under the eyepiece with a 63x objective (Plan Apo 63X/1.4 Oil DIC III). Fluorescent images from the same region of interest were obtained with the excitation-wavelength in the order of 561 nm (for J-aggregates), 514 nm (for JC-1 monomer), and 488 nm (for H2BGFP). Four different regions of interest from the central part of the same retina were imaged before moving to the next retina.

For RH421 (Na^+^/K^+^ ATPase dye) staining, similar steps were taken as for JC-1 staining, with the following modifications: 1) 0.83 μM RH421 dye (61017, Biotium) was added to the glass-bottom culture dishes just before imaging, but not during incubation in the incubator, due to the fast action of RH421. 2) 5 regions of interest were imaged per retina from the central area. 3) The dissection and culture medium were lactate-only medium (see below). 4) Excitation wavelengths: 561 nm (RH421), and 488 nm (H2BGFP).

For imaging genetically-encoded metabolic sensors (PercevalHR, iGlucoSnFR and pHRed), retinas were placed in the incubator for 12 minutes and then taken to confocal imaging without any staining. For the high-glucose condition, the culture medium described above contains ≈15 mM glucose without lactate or pyruvate. For the lactate-only condition, the culture and dissection media were both glucose-pyruvate-free DMEM (A144300, Thermo Fisher Scientific) and were supplemented with 20mM sodium L-lactate (71718, Sigma-Aldrich). For the pyruvate-only condition, the culture and dissection media, were both glucose-pyruvate-free DMEM plus 10 or 20 mM sodium pyruvate (P2256, Sigma-Aldrich). No AAV-H2BGFP was co-injected with these sensors, since the sensors themselves could be used to trace the area of infection. The excitation wavelengths for sensors were 488 nm and 405 nm (PercevalHR, ratiometric high and low ATP:ADP), 488 nm and 561 nm (iGlucoSnFR, glucose-sensing GFP and normalization mRuby), and 561 nm and 458 nm (pHRed, ratiometric low and high pH).

The fluorescent intensity of all acquired images was measured by ImageJ. The ratio of sensors/dye was normalized to averaged control results taken at the same condition.

### Flow cytometry and cell sorting

All flow cytometry and cell sorting were performed on MoFlo Astrios EQ equipment. Retinas were freshly dissected and dissociated using cysteine-activated papain followed by gentle pipetting (Shekhar et al., 2016). Before sorting, all samples were passed through a 35-μm filter with buffer containing Fluorobrite DMEM (A1896701, Thermo Fisher Scientific) and 0.4% BSA. Cones labeled with AAV8-SynP136-H2BGFP (highly cone-specific) were sorted into the appropriate buffer for either ddPCR or RNA-sequencing.

### RNA sequencing

RNA sequencing was done as previously described (Wang et al., 2019). 1,000 H2BGFP-positive cones per retina were sorted into 10 μL of Buffer TCL (Qiagen) containing 1% β-mercaptoethanol and immediately frozen in −80 °C. On the day of sample submission, the frozen cone lysates were thawed on ice and loaded into a 96-well plate for cDNA library synthesis and sequencing. A modified Smart-Seq2 protocol was performed on samples by the Broad Institute Genomics Platform with ∼6 million reads per sample (Picelli et al., 2013). The reads were mapped to the GRCm38.p6 reference genome after quality control measures. Reads assigned to each gene were quantified using featureCounts (Liao et al., 2014). Count data were analyzed using DESeq2 to identify differentially expressed genes, with an adjusted *p* value less than 0.05 considered significant(Anders and Huber, 2010). The raw results have been deposited to Gene Expression Omnibus (accession number GSE161622).

### ddPCR

RNA was isolated from 20,000 sorted cones per retina using RNeasy Micro Kit (Qiagen) as previously described (Wang et al., 2020), and converted to cDNA using the SuperScript III First-Strand Synthesis System (Invitrogen). cDNA from each sample was packaged in droplets for Droplet Digital™ PCR (ddPCR) using QX200 EvaGreen Supermix (#1864034). The reads of expression were normalized to the housekeeping gene *Hprt*. Sequences for RT-PCR primers were designed using the IDT online RealTime qPCR primer design tool. The following primers were selected for the genes of interest: *Txnip* (forward 5’-ACATTATCTCAGGGACTTGCG-3’; reverse 5’-AAGGATGACTTTCTTGGAGCC-3’), *Hprt* (forward 5’-TCAGTCAACGGGGGACATAAA-3’; reverse 5’-GGGGCTGTACTGCTTAACCAG-3’), *mt-Nd4* (forward 5’-AGCTCAATCTGCTTACGCCA-3’; reverse 5’-TGTGAGGCCATGTGCGATTA-3’), *mt-Cytb* (forward 5’-ATTCTACGCTCAATCCCCAAT-3’; reverse 5’-TATGAGATGGAGGCTAGTTGGC-3’), *mt-Co1* (forward 5’-TCTGTTCTGATTCTTTGGGCACC-3’; reverse 5’-CTACTGTGAATATGTGGTGGGCT-3’, *Acsl3* (forward 5’-AACCACGTATCTTCAACACCATC-3’; reverse 5’-AGTCCGGTTTGGAACTGACAG-3’), and *Ftl1* (forward 5’-CCATCTGACCAACCTCCGC-3’; reverse 5’-CGCTCAAAGAGATACTCGCC-3’)).

### Electron microscopy

Intracardial perfusion (4% PFA+1% glutaraldehyde) was performed on ketamine/xylazine (100/10 mg/kg) anesthetized mice before the removal of eyes. The cornea was sliced open and the eye was fixed with a fixative buffer (1.25% formaldehyde+ 2.5 % glutaraldehyde + 0.03% picric acid in 0.1 M sodium cacodylate buffer, pH 7.4) overnight at 4 °C. The cornea, lens and retina were removed before resin embedding, ultrathin sectioning and negative staining at Harvard Medical School Electron Microscopy Core. The detailed methods can be found on the core’s website (https://electron-microscopy.hms.harvard.edu/methods). The stained thin sections were imaged on a conventional transmission electron microscope (JEOL 1200EX) with an AMT 2k CCD camera.

### Statistics

For the comparison of two sample groups, two-tailed unpaired Student’s t test was used to test for the significance of difference, except for P140 *rho*^-/-^ optomotor assay (paired two-tail t-test). For comparison of more than two sample groups, ANOVA and Dunnett’s multiple comparison test was performed in Prism 8 software to determine the significance. A *p* value of less than 0.05 was considered statistically significant. All error bars are presented as mean ± standard deviation, except for the *rd10* optomotor assays (mean ± SEM).

### Study approval

All animal experiments were approved by the IACUC of Harvard University in accordance with institutional guidelines.

## Acknowledgements

We thank Sui Wang, ChangHee Lee, Sylvain Lapan, Gabby Niconchuk, Brian Rabe, Cem Sengel, Sophia Zhao, Yuji Atsuta, Wenjun Xiong, Ryoji Amamoto, Grace Wallick, Gary Yellen, Zhongjie Fu, Zhengping Hu, Maryna Ivanchenko, Paula Montero-Llopis, Microscopy Resources on the North Quad, Maria Ericsson, Electron Microscopy Facility, Flow Cytometry of Immunology, Marcelo Cicconet, Image and Data Analysis Core of Harvard Medical School, Genomics Platform of Broad Institute, Metabolomics Core Resource Laboratory of New York University, Frans Vinberg at University of Utah for discussions and technical support. We also thank Botond Roska, Jacob Keller, Loren Looger, Leah Byrne, John Flannery, William Hahn, David Root, Lewis Cantley, Matthew Vander Heiden, Geoff Wahl, Michael Davidson, Clark Distelhorst, Bong-Kiun Kaang and Thorsten Mascher for plasmids. This work was funded by National Institute of Health (K99EY030951 to Y.X. and U01EY025497 to C.L.C.), Alcon Research Institute (C.L.C.), Astellas Pharmaceuticals (C.L.C.), and Howard Hughes Medical Institute (C.L.C.).

## Author contributions

Y.X. and C.L.C. designed the study and wrote the manuscript with input from other authors. Y.X., S.K.W. and P.R. performed the experiments. Y.X., S.K.W., P.R., E.R.W., C.M.H. and H.F. analyzed the data. D.M.W provided critical software and reagents to this study.

---

**Supplementary Table 1:**
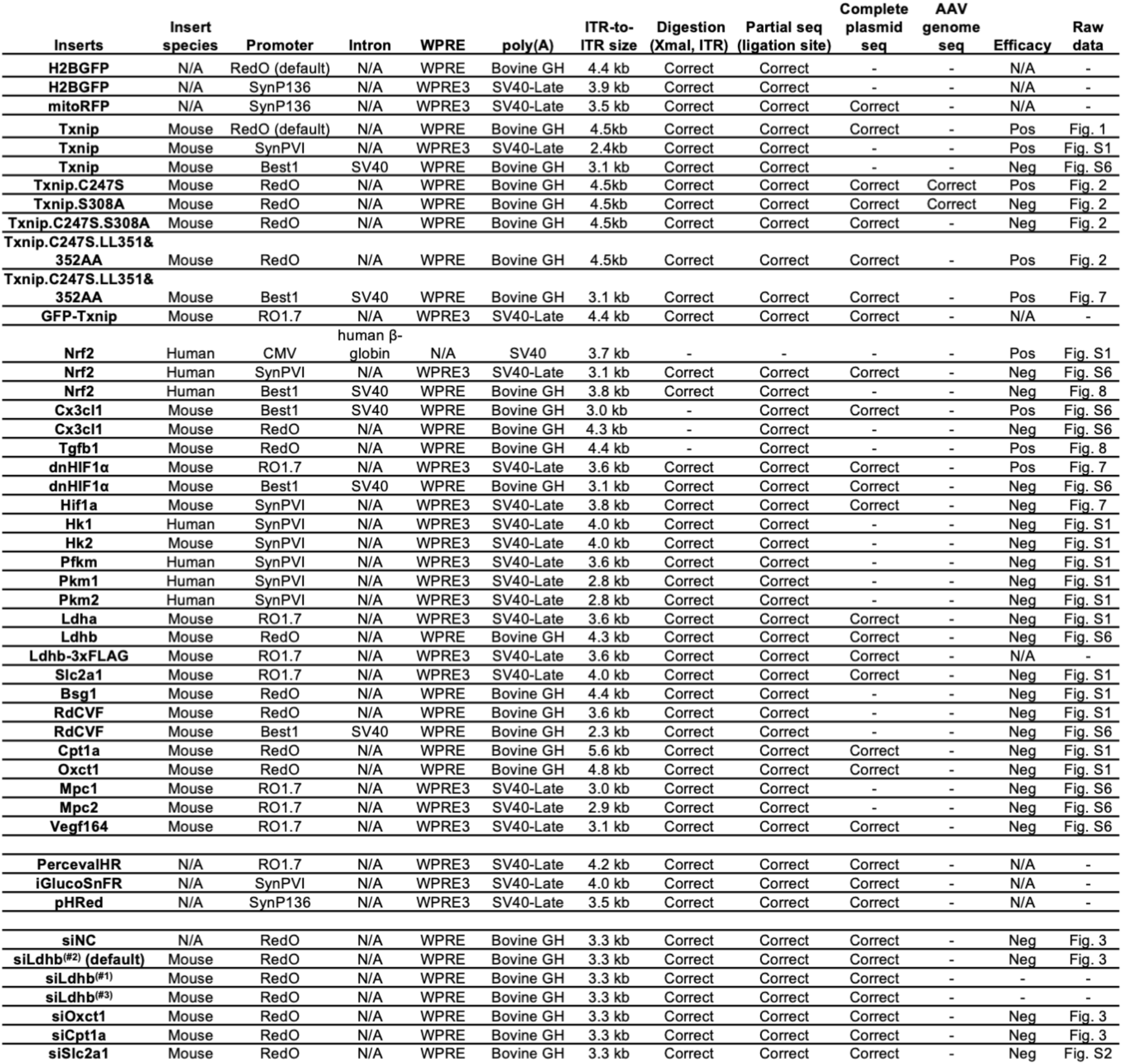
AAV8 vectors used in this study.

**Supplementary Table 2:**
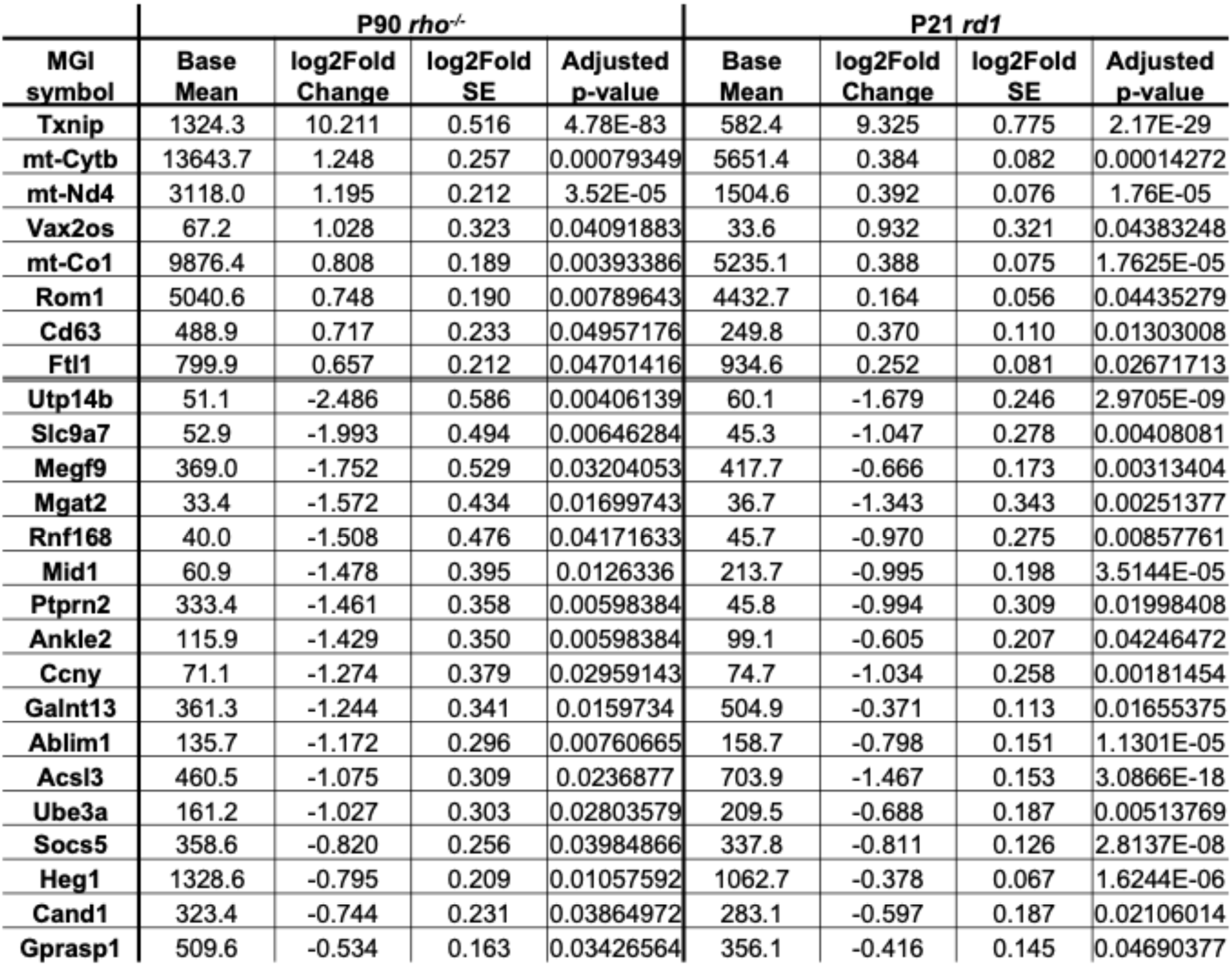
Differentially expressed genes in common between *rd1* and *rho*^-/-^ cones infected by AAV-Txnip.

**Extended Data Fig. 1: Figures related to Fig. 1.**
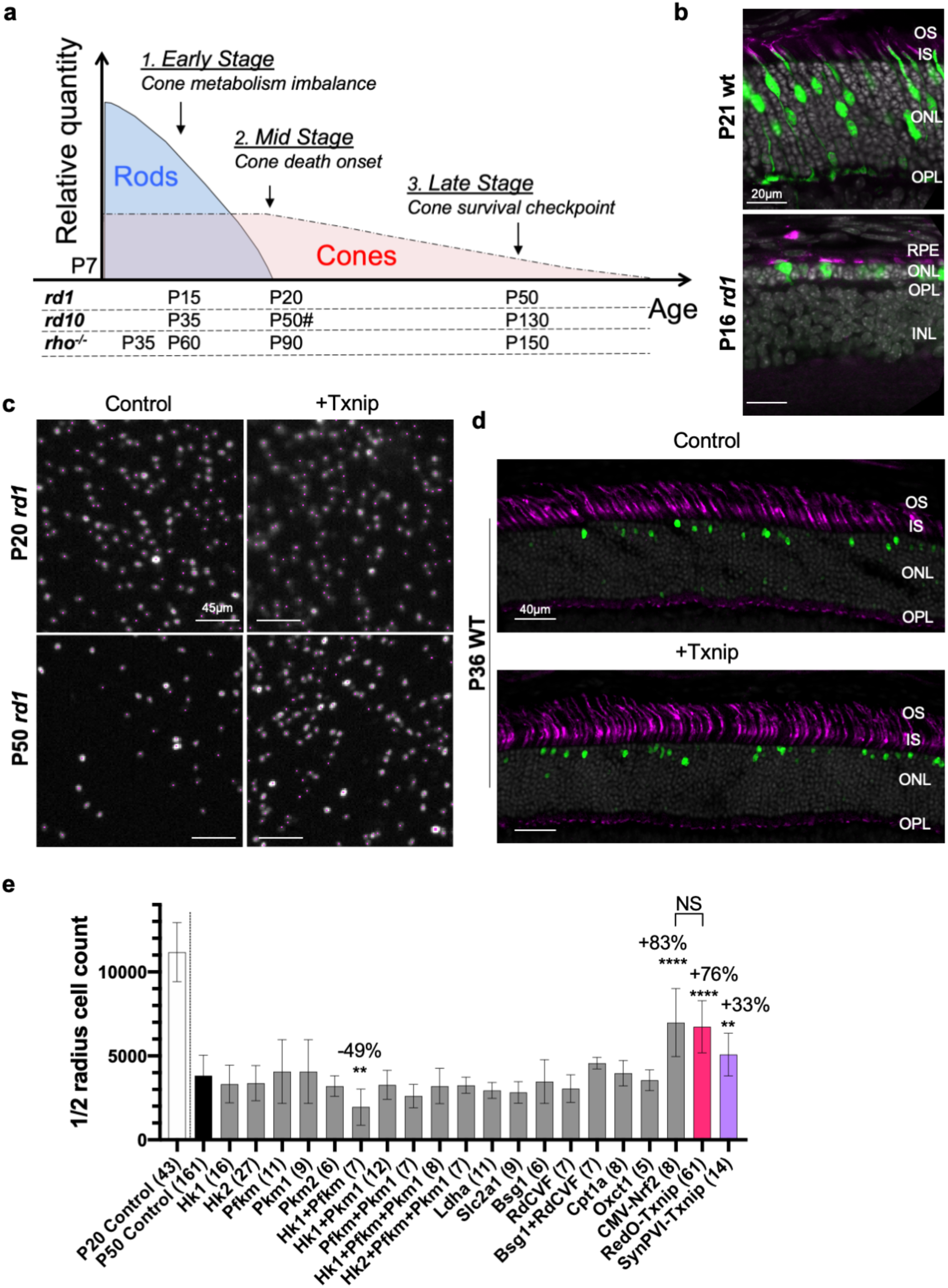
a. Schematics photoreceptor degeneration in RP mice. # *rd10* mid stage varies due to light-dependent rod degeneration (Chang et al., 2007). b. AAV8-RO1.7-GFP.Txnip expression in P21 wt (BALB/c) and P16 *rd1* retina. (Green: GFP. Magenta: PNA for cone extracellular matrix. Gray: DAPI.) Abbreviations. OS: outer segment, IS: inner segment, ONL: outer nuclear layer, OPL: outer plexiform layer, INL: inner nuclear layer, RPE: retinal pigmented epithelium. c. Pixels recognized as cones by a MATLAB automated-counting program zoomed in from the small boxes in the top four panels (Fig. 1a). (Gray: H2BGFP labeled cones. Magenta: center of one labeled cell recognized by MATLAP program.) d. P36 wildtype (C57BL/6J) retinal cross-section with PNA staining injected with control or 2E9 vg/eye RedO-Txnip, indicating RedO-Txnip is not toxic to the wildtype cones. 3E8 vg/eye RedO-H2BGFP was co-injected to track infection. (Magenta: PNA. Green: H2BGFP. Gray: DAPI.) e. Quantification of H2BGFP-positive cones within the half radius of P20 *rd1* control retinas, and P50 *rd1* retinas treated with 20 different vectors and combinations or control. (Please note: we did not use dark-reared *rd10* for testing the RdCVF vector, and our AAV capsid and promoter were different from the original study (Byrne et al., 2015). Error bar: standard deviation. NS: not significant, p > 0.05. * p < 0.05. ** p < 0.01. *** p < 0.001. **** p < or << 0.0001. (Same applies to the rest of Extended Data Figures.)

**Extended Data Fig. 2: Figures related to Fig. 2.**
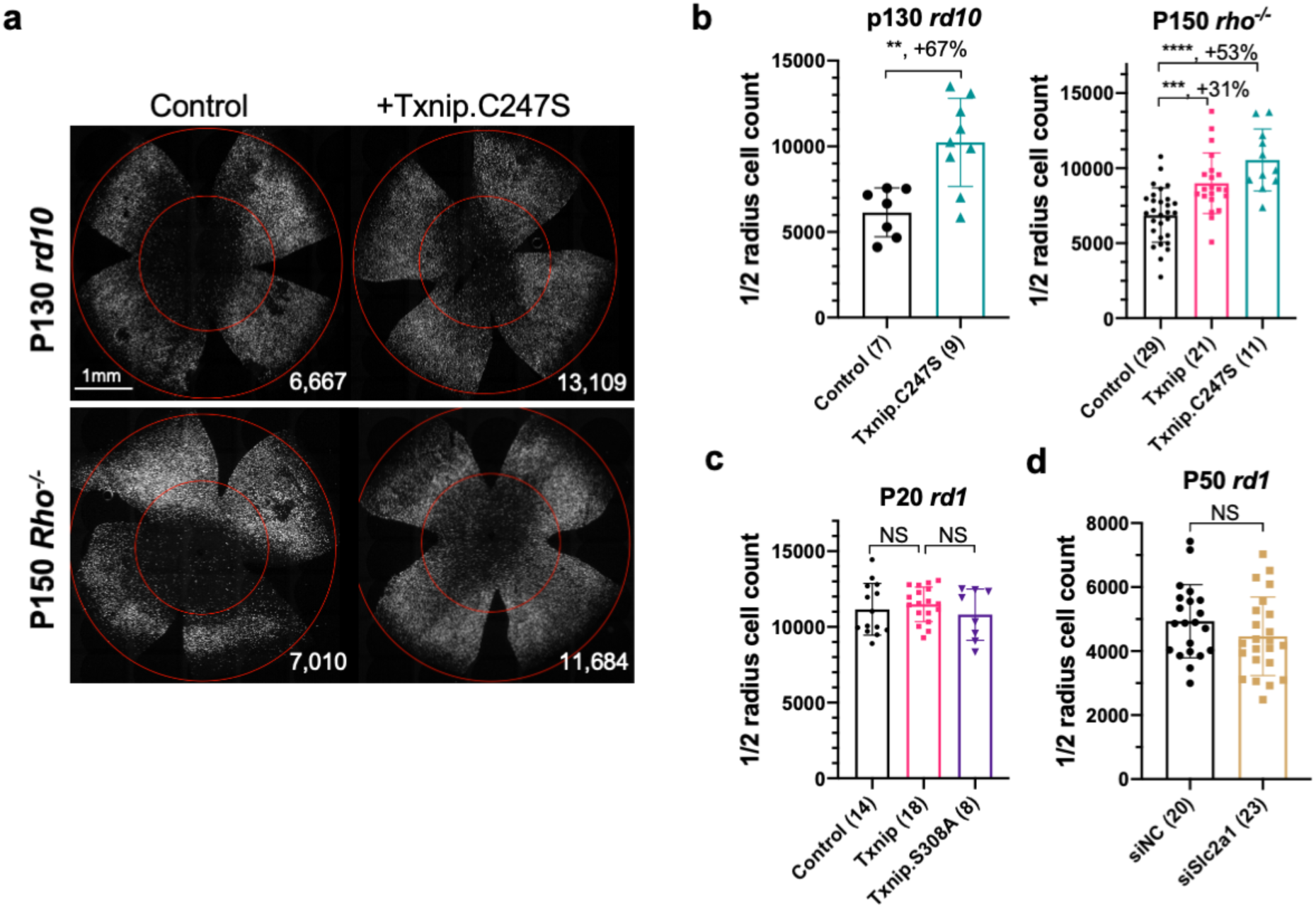
a. Representative P130 *rd10* and P150 *rho*^-/-^ flat-mounted retinas with H2BGFP (gray) labeled cones treated with Txnip.C247S or control. b. Quantification of H2BGFP-positive cones within the half radius of P130 *rd10* and P150 *rho*^-/-^ retinas treated with Txnip.C247S or control. c. Quantification of H2BGFP-positive cones within the half radius of P20 *rd1* retinas treated with Txnip, Txnip.S308A or control. d. Quantification of H2BGFP-positive cones within the half radius of P50 *rd1* retinas treated with siNC (non-targeting scrambled control shRNA) or Slc2a1/Glut1 shRNA.

**Extended Data Fig. 3: Figures related to Fig. 3.**
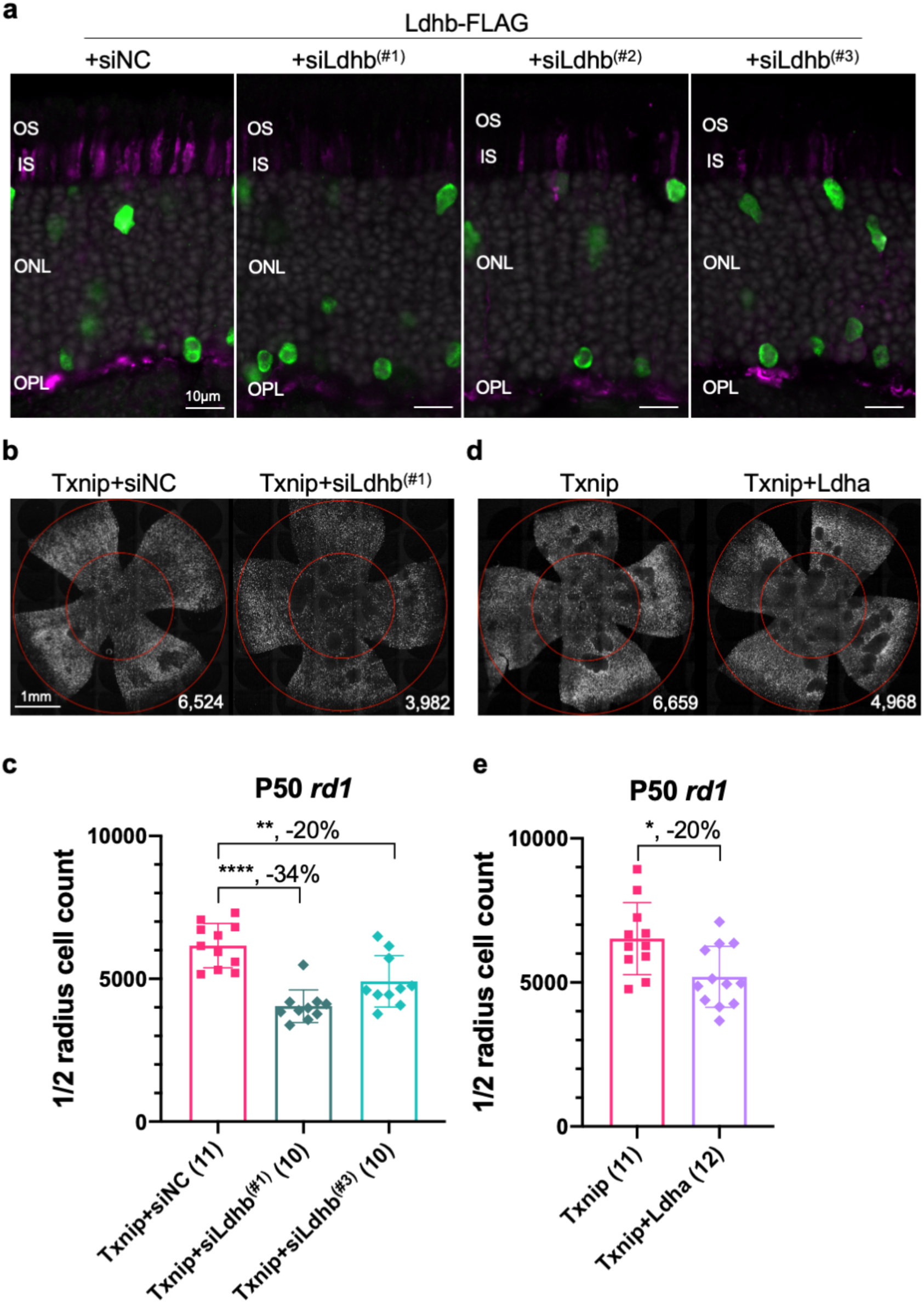
a. AAV8-RO1.7-Ldhb-FLAG with siNC control or *Ldhb* shRNAs in P21 wildtype (CD1) retina plus RedO-H2BGFP to track the infection. (Magenta: anti-FLAG. Green: anti-GFP. Gray: DAPI.) b. Representative P50 *rd1* flat-mounted retinas with H2BGFP (gray) labeled cones treated with Txnip + siNC, Txnip + siLdhb^(#1)^, or Txnip + siLdhb^(#3)^. c. Quantification of H2BGFP-positive cones within the half radius of P50 *rd1* retinas treated with Txnip + siNC, Txnip + siLdhb^(#1)^ or Txnip + siLdhb^(#3)^. d. Representative P50 *rd1* flat-mounted retinas with H2BGFP (gray) labeled cones treated with Txnip or Txnip + Ldha. e. Quantification of H2BGFP-positive cones within the half radius of P50 *rd1* retinas treated with Txnip or Txnip + Ldha.

**Extended Data Fig. 4: Figures related to Fig. 4.**
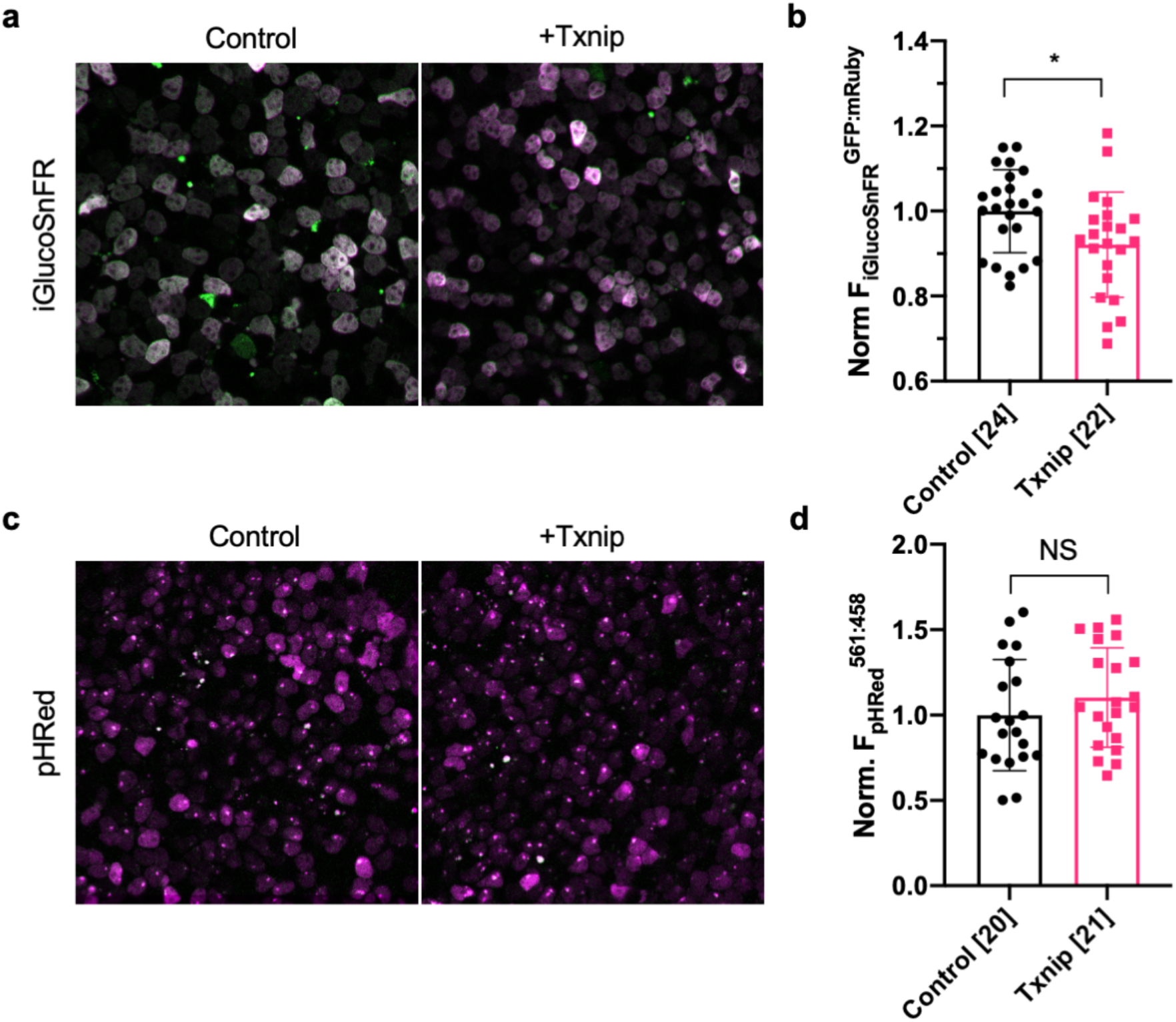
a. Representative *ex vivo* live images of iGlucoSnFR labeled cones in P20 *rd1* retinas cultured with high-glucose medium treated Txnip or control. (Green: glucose sensing GFP. Magenta: mRuby for self-normalization.) b. Quantification of normalized iGlucoSnFR fluorescence intensity (F_iGlucoSnFR_^GFP : mRuby^, proportional to glucose level) in cones from P20 *rd1* retinas treated with Txnip or control (≈3 images per retina). c. *ex vivo* live images of pHRed labeled cones in P20 *rd1* retinas cultured with high-glucose medium treated Txnip or control. (Magenta: fluorescence by 561 nm excitation, indicating a lower pH. Green: fluorescence by 458 nm excitation, indicating a higher pH.) d. Quantification of normalized pHRed fluorescence intensity (F_pHRedx_^561nm : 458nm^, inversely proportional to pH value) in cones from P20 *rd1* retinas treated with Txnip or control (≈3 images per retina).

**Extended Data Fig. 5: Figures related to Fig. 5.**
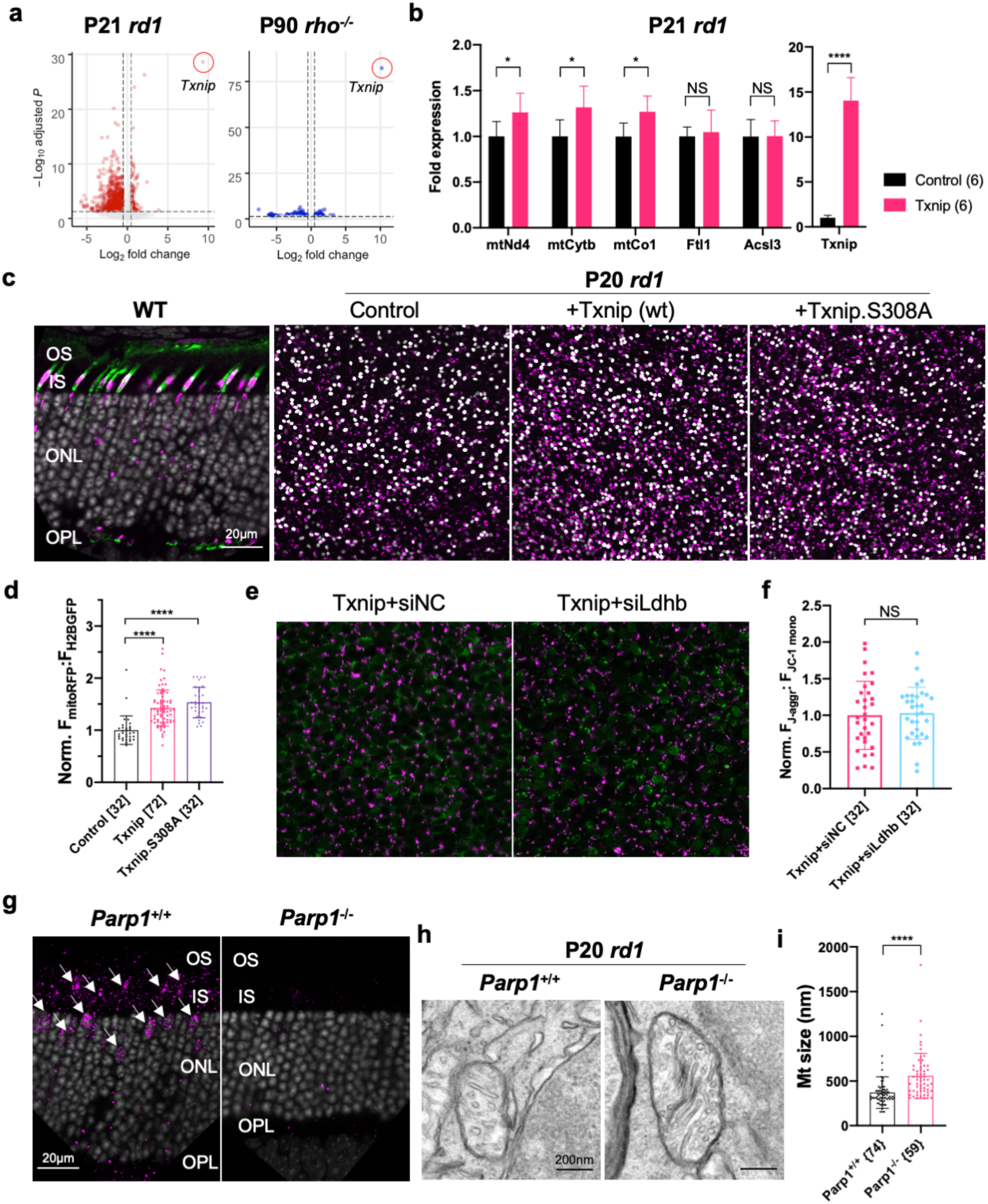
a. Volcano plots of differentially expressed genes in RP cones FACS sorted from P21 *rd1* retinas (left panel; +Txnip, n = 3, relative to control, n = 6) and P90 *rho*^-/-^ retinas (right panel; +Txnip, n = 4, relative to control, n = 4). Dotted lines indicate adjusted *p* < 0.05 and log2 fold change > 0.5. b. ddPCR fold-changes of commonly upregulated mitochondrial ETC genes and genes not confirmed (i.e. *Acsl3* and *Ftl1*) by Txnip overexpression in FACS sorted P21 *rd1* cones. c. First panel: AAV8-SynP136-mitoRFP expression in P26 wildtype (BALB/c) retina cross-section. (Magenta: mitoRFP. Green: PNA. Gray: DAPI.) Other three panels: representative mitoRFP images from the control, Txnip and Txnip.S308A of fixed P20 *rd1* retina flat-mounts near optic nerve head, reflecting the mitochondrial function. (Magenta: mitoRFP. Gray: H2BGFP, for infection normalization.) d. Quantification of normalized mito-RFP:H2BGFP intensity of P20 *parp1* retinas treated with control, Txnip or Txnip.S308A (4 images per retina). e. Representative JC-1 dye staining image from live cones in P20 *rd1* retina treated with Txnip + siNC or + siLdhb^(#2)^. (Magenta: J-aggregate, indicating high ETC function. Green: JC-1 monomer, for self-normalization. H2BGFP channel, the tracer of AAV infected area, is not shown.) f. Quantification of normalized cone JC-1 dye staining (fluorescence intensity of J-aggregate:JC-1 monomer) from live cones in P20 *rd1* retinas treated with Txnip + siNC or siLdhb^(#2)^ (4 images per retina). g. Parp1 antibody staining of *parp1*^+/+^ (C57BL/6J) or *parp1*^-/-^ (on 129S background) retina. (Magenta: Parp1. Gray: DAPI. Arrow heads: Parp1 staining from inner segments and cone nuclei). h. Representative mitochondria EM images from P20 *parp1*^+/+^ or *parp1*^-/-^ *rd1* cones. i. Quantification of mitochondrial diameters from P20 *parp1*^+/+^ or *parp1*^-/-^ *rd1* cones from one retina per condition.

**Extended Data Fig. 6: Figures related to Fig. 7 and 8.**
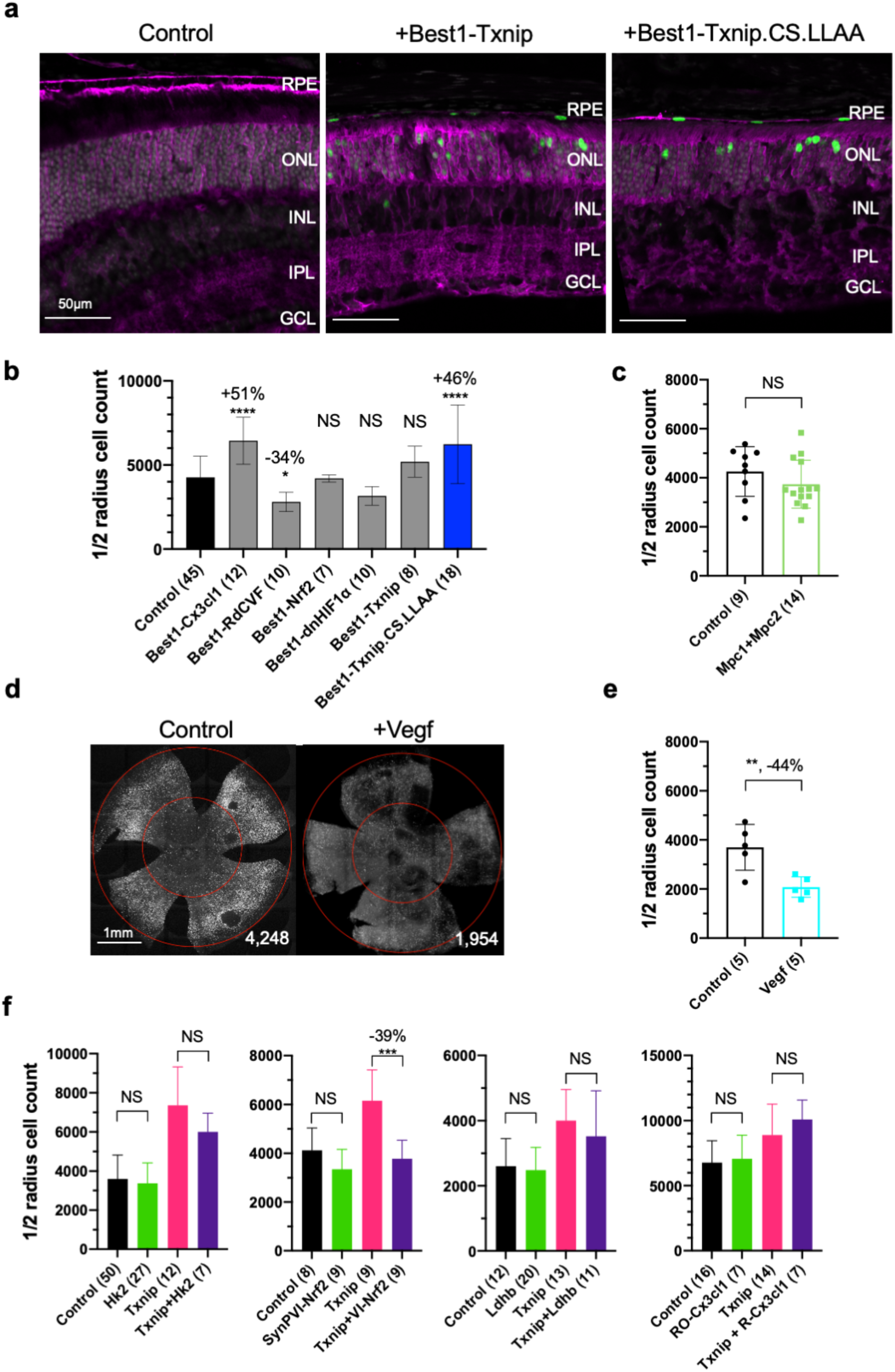
a. Glut1 expression in P37 wildtype (C57BL/6J) eyes treated with control, AAV8-Best1-Txnip or AAV8-Best1-Txnip.C247S.LL351&352AA (Magenta: Glut1. Green: RedO-H2BGFP for infection tracing, leaky expression in RPE due to recombination with Best1-vector due to unclear mechanism(Wu et al., n.d.). Gray: DAPI.) b. Quantification of H2BGFP-positive cones within the half radius of P50 *rd1* retinas treated with 6 different Best1-vectors or control. (Please note: we did not use dark-reared *rd10* for testing the RdCVF vector, and our AAV capsid and promoter were different from the original study (Byrne et al., 2015).) c. Quantification of H2BGFP-positive cones within the half radius of P50 *rd1* retinas treated with Mpc1 + Mpc2 or control. d. Representative P50 *rd1* flat-mounted retinas with H2BGFP (gray) labeled cones treated with Vegf164 and control. Abbreviation: Txnip.CS.LLAA = Txnip.C247S.LL351&352AA. e. Quantification of H2BGFP-positive cones within the half radius of P50 *rd1* retinas treated with Vegf164 or control. f. Quantification of H2BGFP-positive cones within the half radius of P50 *rd1* retinas treated with control, SynPVI-Hk2, SynPVI-Nrf2, RedO-Ldhb, RedO-Cx3cl1, RedO-Txnip and combinations with RedO-Txnip. Abbreviation: VI = SynPVI. RO- or R- = RedO-.

**Extended Data Fig. 7:**
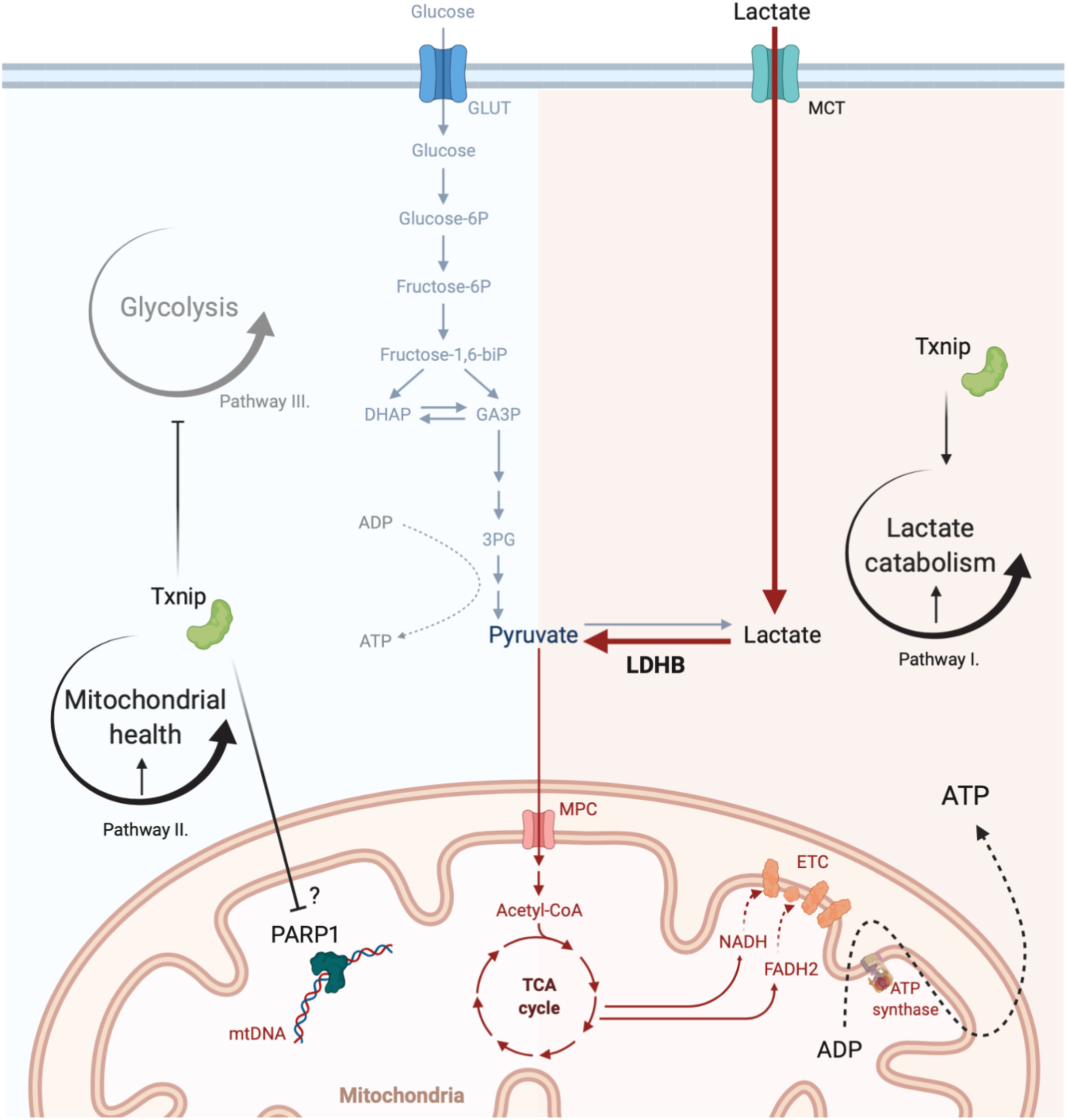
Schematics of proposed Txnip working mechanism.

**Extended Data Fig. 8:**
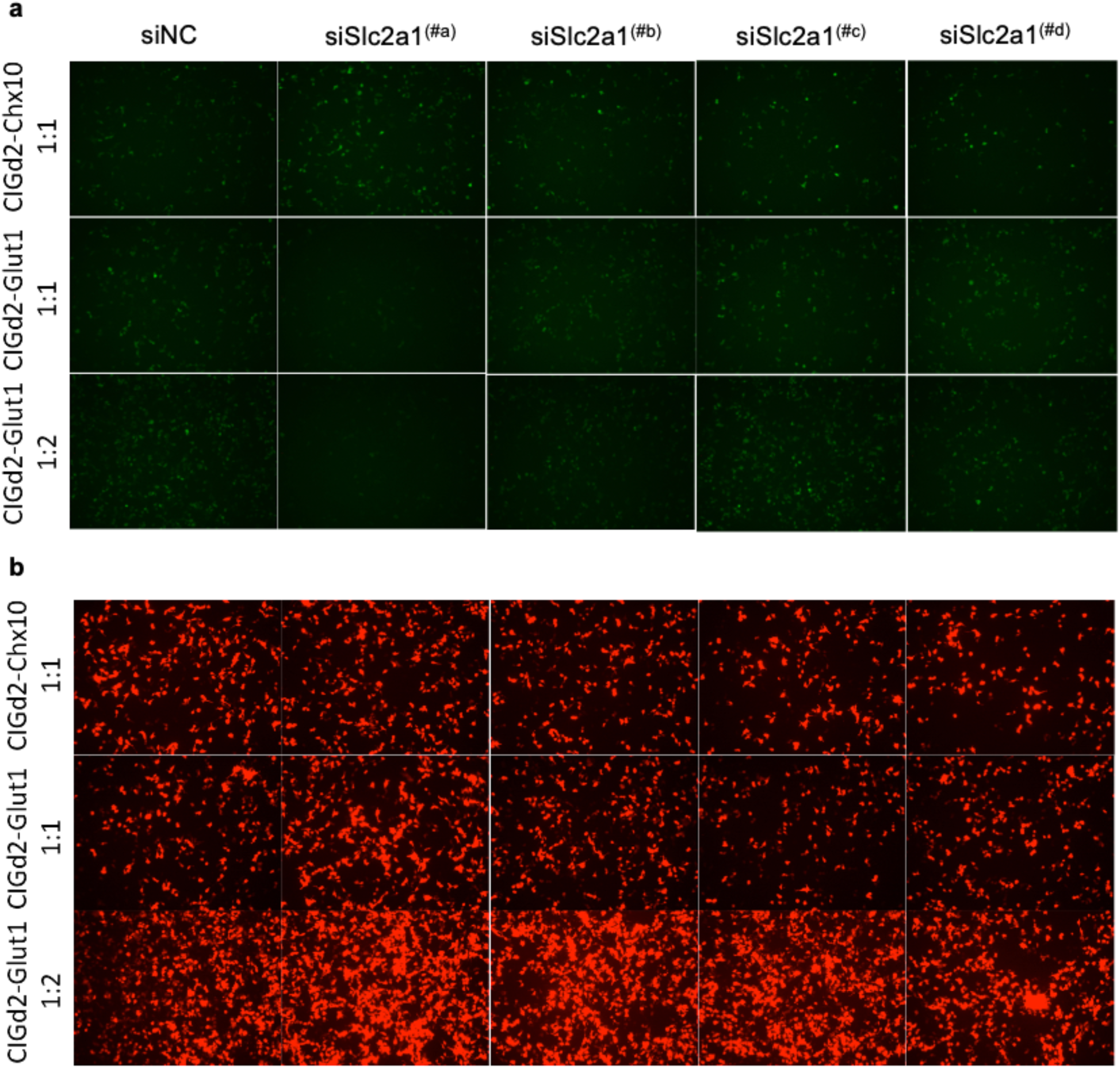
Slc2a1/Glut1 shRNA *in vitro* screening and screened out siSlc2a1^(#a)^ for *in vivo* experiments. a. GFP signals from overnight transfected HEK293T cells labeled with CAG-Slc2a1-IRES-GFPd2 (CIGd2-Glut1) or CIGd2-Chx10 (negative control group) plus siSlc2a1(#a, b, c, d) or siNC at 1:1 or 1:2 ratios. b. mCherry signals (positive-control for transfection) from the same imaging regions as in sub-Fig. a above.

**Extended Data Fig. 9:**
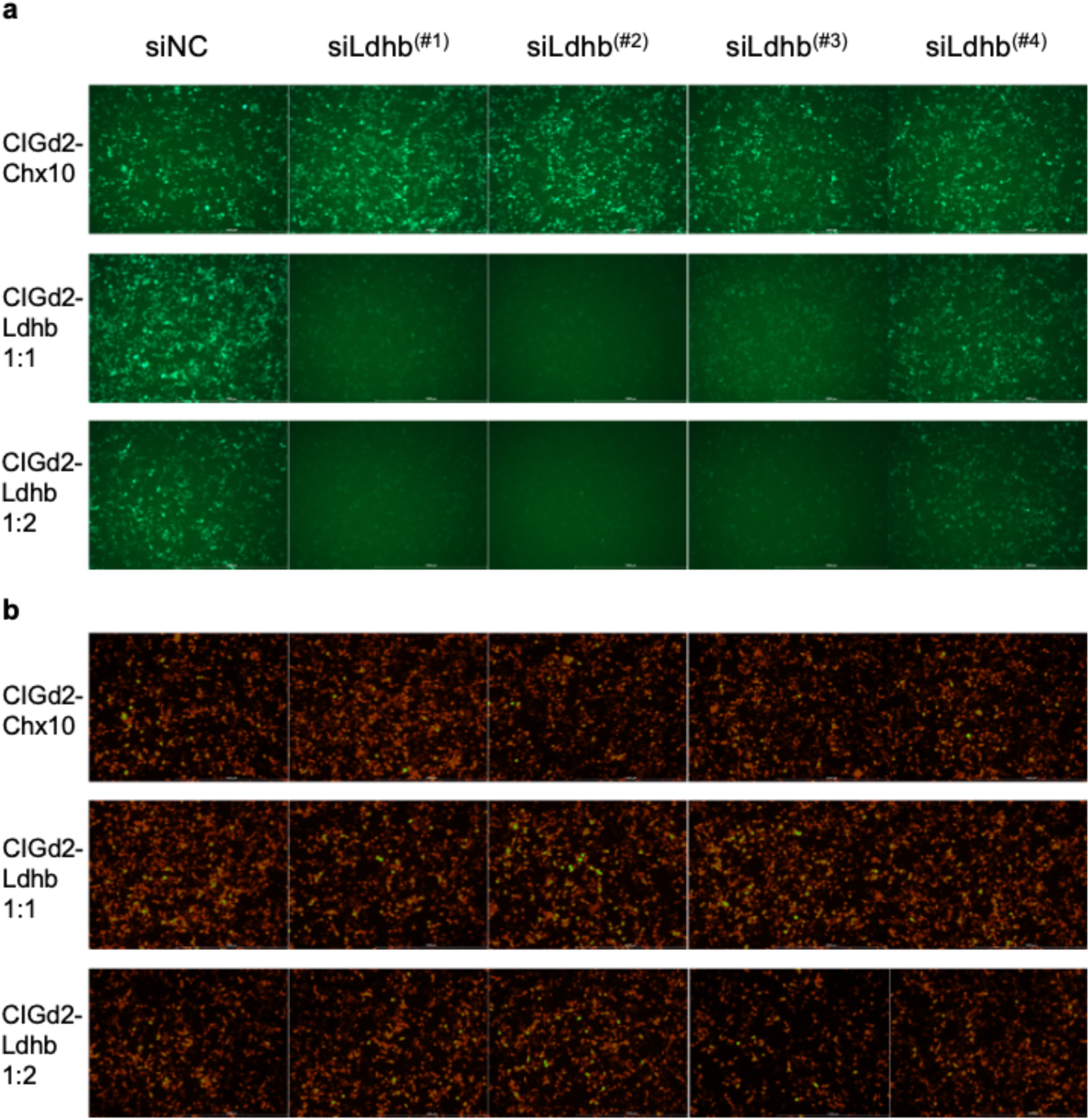
Ldhb shRNA *in vitro* screening and screened out siLdhb^(#2), (#1) & (#3)^ for *in vivo* experiments. a. GFP signals from overnight transfected HEK293T cells labeled with CAG-Ldhb-IRES-GFPd2 (CIGd2-Ldhb) or CIGd2-Chx10 (negative control group) plus siLdhb(#1, 2, 3, 4) or siNC at 1:1 or 1:2 ratios. b. mCherry signals (positive-control for transfection) from the same imaging regions as in sub-Fig. a above.

**Extended Data Fig. 10:**
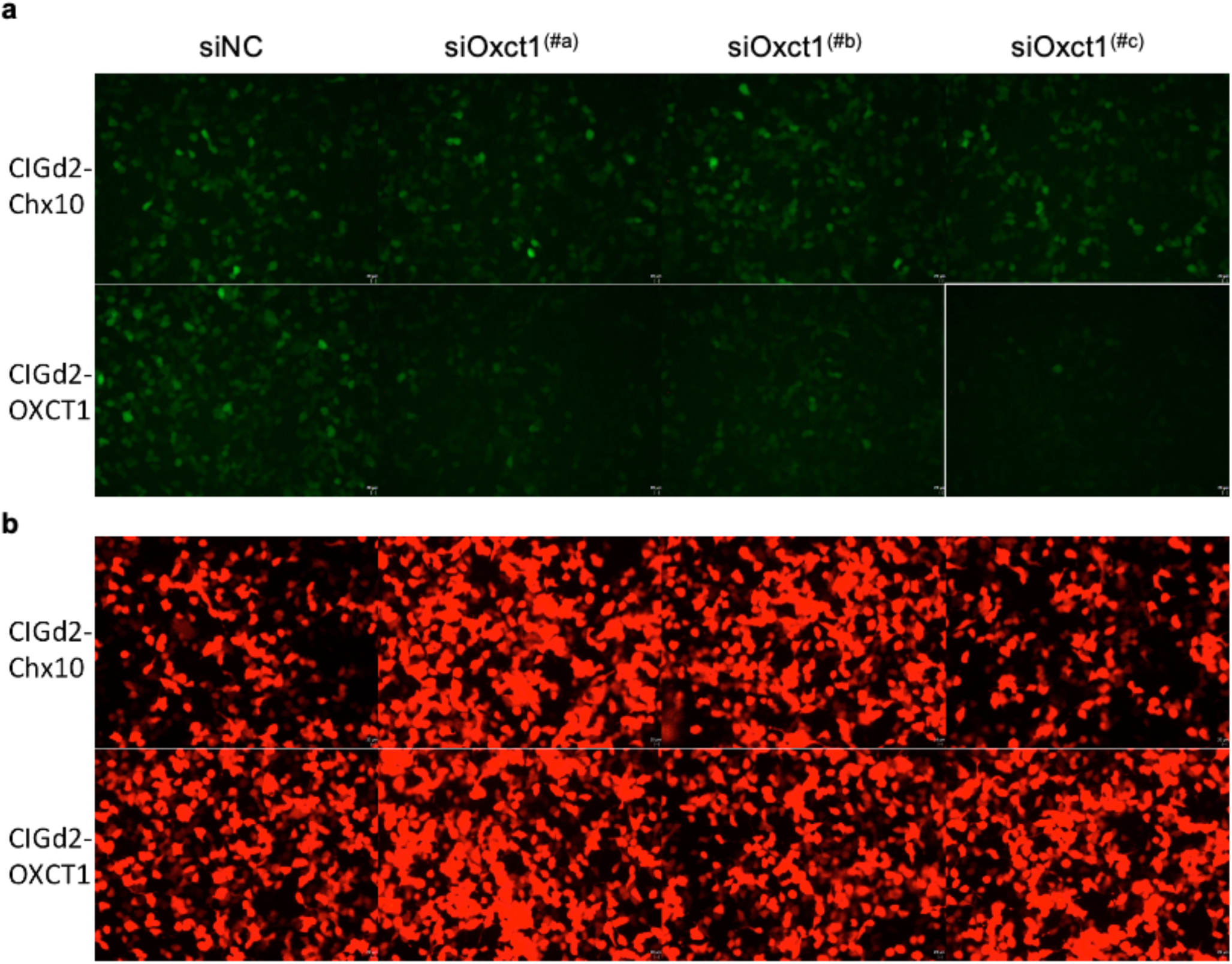
Oxct1 shRNA *in vitro* screening and screened out siOxct1^(#c)^ for *in vivo* experiments. a. GFP signals from overnight transfected HEK293T cells labeled with CAG-Oxct1-IRES-GFPd2 (CIGd2-OXCT1) or CIGd2-Chx10 (negative control group) plus siOxct1(#a, b, c) or siNC at 1:2 ratios. b. mCherry signals (positive-control for transfection) from the same imaging regions as in sub-Fig. a above.

**Extended Data Fig. 11:**
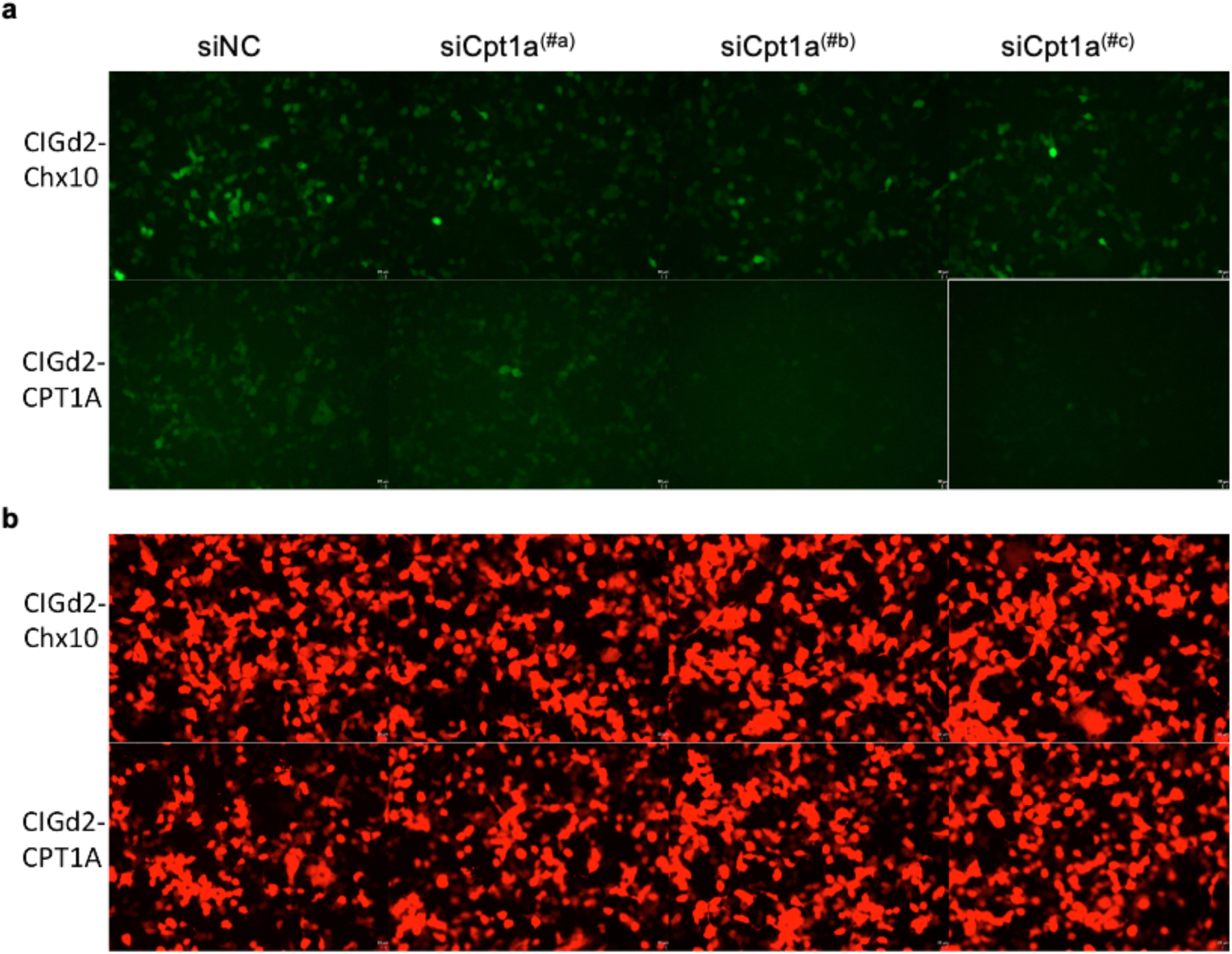
Cpt1a shRNA *in vitro* screening and screened out siCpt1a^(#c)^ for *in vivo* experiments. a. GFP signals from overnight transfected HEK293T cells labeled with CAG-Cpt1a-IRES-GFPd2 (CIGd2-CPT1A) or CIGd2-Chx10 (negative control group) plus siCpt1a(#a, b, c) or siNC at 1:2 ratios. b. mCherry signals (positive-control for transfection) from the same imaging regions as in sub-Fig. a above.

